# Feature and space-based interference with functionally active and passive items in working memory

**DOI:** 10.1101/2024.02.05.578913

**Authors:** Sophia A. Wilhelm, Yuanyuan Weng, Jelmer P. Borst, Robbert Havekes, Elkan G. Akyürek

**Affiliations:** Experimental Psychology, Department of Psychology, University of Groningen; Neurobiology Expertise Group, Groningen Institute of Evolutionary Life Sciences, University of Groningen; Bernoulli Institute for Mathematics, Computer Science and Artificial Intelligence, University of Groningen

**Keywords:** working memory, memory interference, activity-quiescent working memory

## Abstract

Functionally active and passive states in working memory have been related to different neural mechanisms. Memoranda in active states might be maintained by persistent neural firing, whereas memoranda in passive states might be maintained through short-term synaptic plasticity. We reasoned that this might make these items differentially susceptible to interference during maintenance, in particular that passively maintained items might be more robust. To test this hypothesis, we gave our participants a working memory task in which one item was prioritised (active) by always probing it first, while the other item was deprioritised (passive) by always probing it second. In two experiments, on half the trials, we presented an interfering task during memory maintenance, in which the stimuli matched either the feature dimension of the memory items (colour or orientation), or their spatial location. Whether the interfering task appeared on a given trial was unpredictable. In a third experiment where participants were given prior knowledge of the interference condition, and finally in a fourth experiment we used a reward-based prioritisation cue. Across experiments, we found that both active and passive memory items were affected by interference to a similar extent, with overall performance being closely matched in all experiments. We further investigated precision and probability of target response parameters from the standard mixture model, which also showed no differences between states. We conclude that active and passive items, although potentially stored in different neuronal states, do not show differential susceptibility to interference.

**Public significance statement:** The ability to briefly remember information is critical to human cognition. Our so-called working memory is nevertheless rather limited, able to hold only a few items at any one time, and prone to forgetting when we are briefly distracted. Yet, there is reason to believe that not all information in working memory is equally vulnerable. Items that are more passively stored, because they will only be required after some time, might be more resilient to interference. Items that are stored actively, for more immediate recall, might be more easily disrupted. Here, we investigated the effect of an interference task on the retention of both active and passive items in working memory. Our results showed that active and passive items are equally affected by interference, suggesting that resilience in working memory does not depend on the functional state of the items therein.

---

Working memory (WM) is the cognitive system that allows us to store and manipulate information that is not currently available to the senses (Baddeley & Hitch, 1974). Initial evidence of neurophysiological studies on working memory have proposed ongoing neuronal activity as a mechanism of storing information in WM (Goldman-Rakic, 1995). However, in recent years this view that the underlying mechanism of WM is exclusively dependent on ongoing neuronal firing has been questioned (Stokes, 2015). Indeed, intact WM behaviour can be observed even in the absence of persistent neuronal firing (LaRocque et al., 2012; Muhle-Karbe et al., 2021; Rose et al., 2016; Wolff et al., 2017), and even in situations where an interference task was presented to disrupt ongoing (alpha) activity (Oberauer & Awh, 2022). Thus, evidence is accumulating for maintenance mechanisms of working memory that do not exclusively rely on persistent firing. Rather, there also seems to be an activity-silent, or at least activity-quiescent, state that can support the maintenance of working memory content (Stokes et al., 2015).

Short-term synaptic plasticity (STSP) has been proposed as a potential mechanism underlying the maintenance of quiescent WM traces (Kozachkov et al., 2022; Miller et al., 2018; Stokes, 2015). According to this view, after initial encoding items could be stored in rapid changes in synaptic weights within an assembly of neurons. Computational evidence supports this hypothesis: Residual calcium from the initial encoding phase of WM can produce synaptic traces via STSP that can be read out later (Mongillo et al., 2008). Similarly, the inclusion of such STSP mechanisms in an artificial recurrent neural network seems to be necessary to produce firing patterns like those observed in primates during WM maintenance, as well as to attain robustness against network degradation (Kozachkov et al., 2022). Finally, the patterns of working memory maintenance previously seen in EEG recordings of human participants can be reproduced with a functional spiking-neuron model by means of a calcium-based STSP implementation (Pals et al., 2020). Collectively, these findings suggest that STSP provides a plausible mechanism for activity-quiescent WM states.

It seems reasonable to assume that both persistent-activity and activity-quiescent states support working memory, because they serve different functional roles. Persistent-activity states have been associated with functionally active items, that is, items maintained in the focus of attention and/or prioritised items needed for immediate recall. Activity-quiescent states have in turn been associated with functionally passive items; items which are currently deprioritised, outside the focus of attention and only needed for later recall. For example, in a classic two-item retro-cueing paradigm, an item that is known to be relevant only later in the trial (i.e., deprioritised) first enters an activity-quiescent state, but is transformed back into a persistent-activity state once a response to the prioritised item has been made, and a response to the formerly deprioritised item will soon be needed (LaRocque et al., 2012; Wolff et al., 2017). Furthermore, items can be moved back adaptively between persistent-activity and activity-quiescent states on a single-trial basis, based on which item was cued to be relevant on the current trial (Muhle-Karbe et al., 2021). The existence of persistent-activity and activity-quiescent states as two functional storage systems within working memory prompts the question of what the advantages are of having different ways to store items in WM in the brain, and in which situation each mechanism might come into play. A quiescent state of WM maintenance could be more energy-efficient (Stokes, 2015), but one could also wonder whether there could be functional benefits of having distinct neuronal states which can support working memory maintenance. One hypothesis is that quiescent states might offer greater protection against interference from incoming sensory information, as synaptic traces are less dynamic and therefore potentially less vulnerable than ongoing neuronal firing (Kozachkov et al., 2022; Lundqvist, Herman, Warden, et al., 2018; Miller et al., 2018). This is what we set out to test in the present study.

The susceptibility of prioritised versus deprioritised items to interference within WM is unclear. Depending on the study, prioritised items may be protected from interference (Makovski et al., 2008; Makovski & Pertzov, 2015; Schneider et al., 2017; Souza & Oberauer, 2016), may be more susceptible to interference (Hitch et al., 2018; Hu et al., 2014; Mallett & Lewis-Peacock, 2019), so similarly affected by interference (Zhang & Lewis-Peacock, 2023). Some of this ambiguity might have arisen because interference effects in WM are influenced by both the nature of the distractors and the mode of prioritisation of memory items. Factors such as featural similarity and spatial overlap between distractors and memory representations affect the extent of disruption (Hu et al., 2014; Pinto et al., 2013). For example, suffix tasks or visual distractions have been shown to disrupt WM representations only when distractors overlap with memory features (Allen et al., 2015; Allen & Ueno, 2018; Hu et al., 2014). Furthermore, while prioritisation through retro-cueing or reward-driven selection enhances the accessibility of prioritised items, it does not necessarily confer immunity against interference (Zhang & Lewis-Peacock, 2023).

In the current set of experiments, we employed a block design where memory items were explicitly prioritised (functionally active) or deprioritised (functionally passive). A distractor task, involving visual discrimination, was presented during the maintenance phase, and the interference was systematically manipulated to overlap with the active or passive items. Importantly, these interference manipulations included either featural (color or orientation) or spatial overlap, as low-level sensory representations in WM might encode spatial attributes, even when not explicitly relevant to the task. This approach allows us to make a comparison between feature and space-based interference, and their potentially differential effects on active versus passive states. We also included an experiment in which participants were cued about upcoming interference, to assess the ability of our participants’ to strategically shield active or passive memory representations. Anticipation and strategic adjustments can impact WM robustness (Banks et al., 2023; Gresch et al., 2021; Klink et al., 2017), yet whether these effects differentially apply to active versus passive states remains unclear.

To preview the results, we consistently found that both active and passive items were susceptible to interference, with no evidence that a passive and potentially quiescent neuronal state provided a different (higher) level of protection against interference. Interference effects were driven by reductions in precision and probability of target responses for both states, and these effects were stronger when the interfering stimulus matched the memory item, regardless of priority order, and regardless of whether participants had explicit knowledge of the interference feature or location, or not. Interference effects thus appear to be robust across memory states and neither active nor passive memory states provide increased protection from interference.

## Method

### Participants

Across all four experiments, 205 participants (mean age = 20.9, SD = 3.6; 159 female) were either recruited as first year psychology students participating in the experiments for course credit, or through the paid-participant pool of the department where they received monetary reward in exchange for participation. In experiment 1, we collected data from 51 participants. For experiment 2, we collected data from 43 participants, for experiment 3 we collected data from 51 participants, and for experiment 4 we collected data from 60 participants. We are interested in fundamental aspects of basic cognition and therefore believe that using young-healthy adults provides an appropriate sample. However, it should be noted that we only sampled from first-year psychology students, which means our sample was also highly educated, and which might limit the translation to other groups within society. It should also be noted that our sample was largely female.

As we performed all analyses using Bayesian statistics, we opted for optional stopping to determine our sample size. We collected data until we found sufficient evidence for either the alternative or the null hypothesis. We aimed to find at least very strong evidence for the interference effect based on absolute error (according to Wetzels at al., (2011) BF_Alternative_/BF_0_ 30-100). As we performed optional stopping based on the interference effect of absolute error, some of our Bayes factors for the model parameters were lower, and produced only anecdotal evidence (according to Wetzels et al. (2011) BF_Alternative_/BF_0_ = 1-3). For these model parameters we thus cannot conclude an effect or a lack thereof for certain. The study was conducted in accordance with the Declaration of Helsinki (2008) and was approved by the Ethical Committee of the Faculty of Behavioural and Social Sciences at the University of Groningen (Study Code = PSY-2122-S-0206).

### Apparatus and stimuli

The experiment was programmed using OpenSesame, a freely available software tool for designing experiments (Mathôt et al., 2012). Stimuli were presented using a 19-inch CRT screen, at a resolution of 1,280 x 1,024 pixels with a refresh rate of 100Hz. Participants were seated approximately 60 cm away from the screen in a sound-attenuated testing chamber with dimmed lights. A grey background (RGB: 128) was maintained throughout the experiment. The experimental stimuli covered a circular area of 6.69° degrees of visual angle on the screen. Orientation gratings were presented at a contrast of 80% relative to the background, with a spatial frequency of 0.05 cycles per degree. Colours were presented on the CIElab colour space (L*=70, a = 0, b = 0, radius = 38).

### Experiment 1

In Experiment 1 (N= 51), we wanted to test if memory items in an active (prioritised) state were more prone to feature-based interference compared to items in a passive (deprioritised) state. At the start of the experiment, participants were given written and verbal instructions and signed written informed consent. Participants started with several practice trials, after which they were given the opportunity to ask questions before moving on to the experimental trials.

The experiment was divided into blocks. Each block started with a block cue, either stating ‘colour’ or ‘orientation’ which was indicative of the response order for the two memory items at the end of each trial. On any given trial, participants were always presented with one colour as a memory item and one orientation as a memory item (Figure 1A). The presentation order of the memory items was counterbalanced but randomized and independent from the block cue; thus, the block cue did not indicate if the colour or orientation memory item was presented first but was purely informative for response order at the end of the trial. The block cue was only shown before the first trial of each block, participants then did 24 trials.

**Figure 1.**
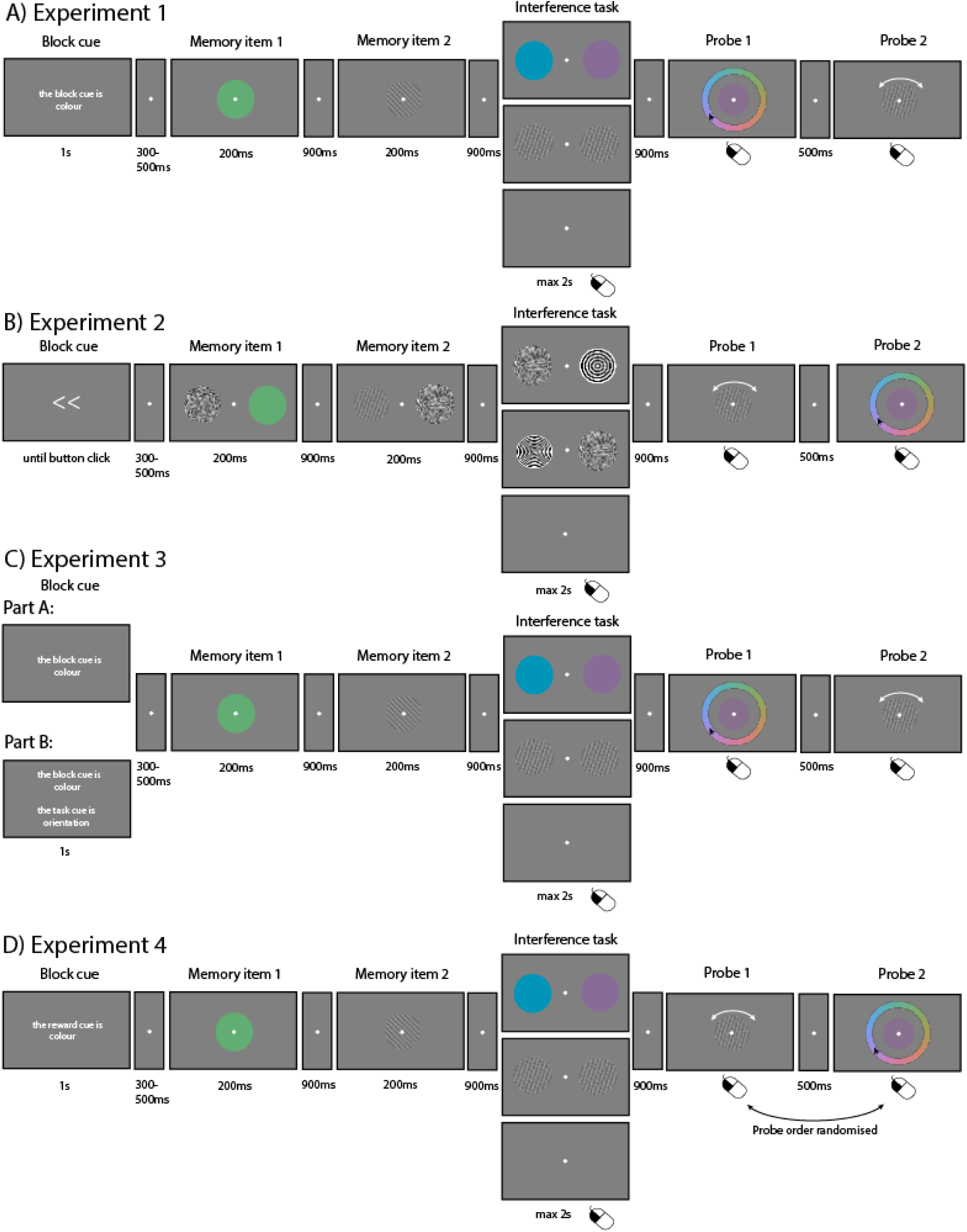
Overview of the trial structure per experiment. *Note:* Overview of the trial structure for each experiment. **A**: Experiment 1, **B**: Experiment 2, **C**: Experiment 3, **D**: Experiment 4.

Each trial consisted of an initial fixation dot between 300-500ms (uniformly sampled), followed by the presentation of the first memory item for 200ms. After a fixation dot of 900ms, the second memory item was presented for 200ms. Another fixation dot was presented for 900ms, after which the interference task was presented. For the interference task participants were presented with two items laterally on the screen at the same time. The interference items could either consist of two coloured circles or two Gabor patches. The task was to judge if the two items presented on the screen were the same (i.e., the same colour or same orientation) or different from each other. The two items were the same for 50% of the trials, and different for the other 50% of trials. We drew colours and orientations randomly from the full circle; thus between 1-180 degrees for orientations and 1-360° degrees for colours. Participants indicated their response by using the different mouse buttons to indicate if the items presented were the same or different from each other (left and right mouse buttons representing same or different conditions was counterbalanced between odd and even participant numbers). Participants had up to 2000ms to give a response, until the experiment would move on. We also included a baseline condition where instead of the interference display, the fixation dot stayed on the screen for 2000ms and participants were instructed to do nothing but to keep looking at the screen. After a response was given in the interference conditions, a fixation dot was presented for 900ms, after which the response screen for memory item 1 appeared, followed by the response screen for memory item 2. Participants could give their response to memory items by selecting the correct colour from a colour wheel for the coloured memory item, and by rotating a Gabor patch in the correct orientation that matched the orientation memory item. The response screens for response 1 and response 2 were separated by a fixation dot, which was presented for 900ms. Participants’ response time to respond to the memory items was not limited.

The logic behind the interference manipulation was as follows: On a given trial, if the feature dimension of the active item matched with that of the interference task, we call this a match trial for the active item. If the feature dimension of the active item did not match with the feature dimension of the interference task; we call this a no match trial for the active item. The same logic applies to the passive item. In both cases, a match between interference task and memory item should lead to higher interference than the absence thereof.

Participants completed 24 practice trials followed by 384 experimental trials divided into 16 blocks, which took approximately 75 minutes to complete. Practice trials were excluded from any analysis. Each interference condition occurred equally often, such that there were 128 trials of each condition per participant. All manipulations were randomised and counterbalanced.

### Experiment 2

The research question of Experiment 2 (N=43) was identical to Experiment 1, but rather than interfering with memory items at the featural level, the manipulations were now space-specific. Unless stated otherwise, the procedure was exactly the same as in Experiment 1. Again, we used a block design where we now cued the location of the item that would be probed first at the end of the trial. On any given trial, the participant would then see two memory items presented laterally at an equal distance of 5.93 degree of visual angle from the fixation dot (Figure 1B). The memory items were presented one at a time with a noise patch being presented in the empty location. The noise patch consisted of a Gaussian noise patch. The memory items were identical to Experiment 1 and the order in which they were presented was randomised and counterbalanced independent from the block cue. The interference task was presented at the same locations as the memory items. For the interference task, in one location the participant would see a black-and-white ‘bull’s eye’ stimulus, and the task was to judge if the spatial frequency of this item was high or low. The corresponding spatial frequencies were 0.5 (low spatial frequency) and 1.4 cycles per degree (high spatial frequency). The ‘empty’ location would be filled with a noise patch again identical to the ones used during memory item presentation. Mouse buttons were used to respond to the interference task where one mouse button would correspond to high spatial frequency and the other would correspond to low spatial frequency (mouse buttons were counterbalanced between even and odd participants). At the end of the trial, the participant had to respond to both memory items in the order indicated by the block cue.

As in the previous experiment, the logic of interference or no interference conditions remained the same but was now not dictated by a match of feature dimension between a memory item and the interference task but rather by a match or no match of its spatial location with the interference item to make it a match or no match trial, respectively.

Participants completed 24 practice trials followed by 384 experimental trials divided into 16 blocks, which took approximately 75 minutes to complete. Practice trials were excluded from any analysis. As before, each interference condition occurred equally often, such that there were 128 trials of each condition per participant. All manipulations were randomised and counterbalanced.

### Experiment 3

In experiment 3 (N=51) we wanted to test whether the predictability of the interference feature would influence the magnitude of the interference effect. This experiment had two phases. In the first part, we replicated the exact set-up of experiment 1. Participants did 180 trials in 10 blocks in this condition. Once they finished these, the participants moved onto the second part. In this part, we kept the previous block cue; however, we also blocked the interference condition and explicitly cued its feature dimension as well. Thus, participants had now prior, explicit knowledge of the interference condition. In this condition, participants performed 192 trials across 12 blocks. Participants performed 24 practice trials of part 1 which were not part of any analysis to familiarise themselves with the task. As in the previous experiments, within the blocks of each conditions, the three interference trials were equally divided between the trials. On average it took participants 90 minutes to complete the experiment. All manipulations were randomised and counterbalanced.

### Experiment 4

Due to the design of the experiments, in the three previous experiments the prioritisation manipulation was confounded by response order, as the prioritised item was always tested first. In experiment 4 (N = 60) we therefore changed the prioritisation approach. We added a reward manipulation that gave participants higher points for responding correctly to the cued (prioritised) item. Unless stated otherwise, the procedure closely followed the design of Experiment 1. As previously, each block started with a cue stating either ‘colour’ or ‘orientation’. However, rather than indicating the response order, the cue now indicated which feature dimension would be more strongly rewarded for a correct response in that block. For a correct response to the cued item participants received three points, whereas the uncued item was worth just one point. Participants were accumulating points throughout the trial, and this was translated into money at the end of the experiment. The maximum reward value was set to 8€. On any given trial, participants were again presented with one colour and one orientation as memory items in randomised order, followed by the same interference task and subsequent response to both memory items. In contrast to Experiment 1, the order in which colour and orientation memory items were probed at the end of the trial was now randomised and counterbalanced and thus independent of block cue. Participants completed 24 practice trials followed by 288 experimental trials divided into 10 blocks which took approximately 75 minutes to complete. Practice trials were excluded from any analysis. All other aspects of the procedure were identical to Experiment 1. All manipulations were randomised and counterbalanced.

### Analysis

The data of all three experiments were analysed using the same approach. Participants whose response error was +/-2SD larger than the mean, or participants who performed (below) chance level in the interference task were excluded from the analysis (Exp 1: N=3, Exp 2: N=2, Exp 3: N=7, Exp4: N=4).

To quantify memory performance of participants we calculated the absolute error response for each trial for each memory item type (active versus passive), and each condition (match, no match and baseline). To render the two feature-spaces directly comparable, absolute orientation errors (originally on a 0–180° scale) were multiplied by two, thereby mapping them onto the same 0–360° range as the colour errors. To test for differences between conditions on each probe we used separate Bayesian repeated-measures ANOVAs to investigate if there was a main effect of interference condition. In cases where we found a main effect of condition, we used paired-samples Bayesian t-tests, which allowed us to quantify evidence for an interference effect (H_A_), as well as to quantify evidence for no effect of interference (H_0_).

To quantify if there were any differences in magnitude between the interference effect on active versus passive items we calculated z-scores and statistically compared those. We chose to compare z-scored responses rather than absolute error since the absolute error for the passive item was on average higher. Thus, we z-scored all individual absolute error responses and calculated an average z-score for the interference and no interference condition per participant. Again, we used Bayesian t-tests to test for statistical differences in the magnitude of interference.

We furthermore wanted to quantify the nature of interference by fitting the two-component standard mixture model (Zhang & Luck, 2009). We fitted independent models for both active and passive items for each condition using the mixture package (Grange & Moore, 2022; version: 1.2.1). The standard mixture model provides two parameters based on the error distribution. It provides the probability of an item being stored in memory (*P_m_*), with smaller values of *P_m_* indicating an increase in guessing responses. As a second parameter it provides the precision (*k*) of the memory representation. Relating this to our experiments, this provided us the opportunity to test if interference leads to a gradual imprecision of the memory representation, which would be reflected in an effect on *k*, or if interference leads to a complete loss of the memory representation, which would be reflected in an effect on *P_m_*. To test statistical differences between the parameters we used Bayesian repeated measures ANOVAs followed by paired-sample Bayesian t-tests.

Pre-processing, mixture modelling, and plotting of the data was performed using R (R Core Team, 2018; version 4.3.2). Bayesian statistics were calculated using JASP (JASP Team, 2023; version: 0.18.1), and Bayesian values were interpreted based on Wetzels et al. (2011). Bayesian statistics in JASP were calculated using the default priors provided in the software. For the Bayesian repeated-measures ANOVAs the prior is a uniform Cauchy distribution – which assumes that all models are equally likely - and set to 0.5 and 1 for fixed and random effects, respectively. For the paired-sample t-test the default prior is also based on a Cauchy distribution and is set to 0.707. We performed a robustness check, as given in JASP, which systematically changes the Cauchy prior and tests how this influences the Bayes factor. We performed this check for all post-hoc t-tests done on the absolute error analysis and modelling parameters. We found that our Bayes factors for effects that had at least substantial evidence (BF_10_>3) were robust and showed qualitatively similar results. In some cases where our model parameters were only supported by anecdotal evidence, the robustness check showed qualitatively different results in the case of extreme priors (Cauchy prior of 1.4, or max prior to obtain evidence in favour of H_A_). Overall, we concluded that our results showed robust results, as only extreme priors for anecdotal evidence showed qualitatively different results; and as anecdotal evidence is by definition not definitive these observations are to be expected.

The raw datasets and datasets used to perform the statistical analysis in JASP can be accessed here (this link will be made public upon publication): https://osf.io/qpt5f/?view_only=644630a919ff4394905a20e7a296fb89

### Transparency and openness

In the above, we reported how we determine our sample size, criteria for data exclusions and all experimental manipulations. Experiments were performed in the same order as reported in the manuscript and all data was collected in 2024, except for Experiment 4 which was collected in 2025. Cleaned datasets, modelling outputs, and datasets used for statistical analysis in JASP are available through the link above. The experiments in this paper and their analyses were not preregistered.

## Results

### Experiment 1

In Experiment 1 (N=48), we tested whether feature-based interference differed between active and passive items held in working memory. The results are shown in Figure 2. Across interference conditions, we found decisive evidence for better performance on active items (μ_AS_ = 18.6°, sd_AS_ = 6.6°), compared to passive items (μ_AS_ = 22.8°, sd_AS_ = 8.0°) (*t_paired_* BF_10_ > 100). Furthermore, we observed decisive evidence that participants performed above chance level for the interference task (μ = 89.4%, sd = 5%) (*t_one-sided_* BF_10_ > 100), indicating that participants paid attention to the stimuli.

**Figure 2.**
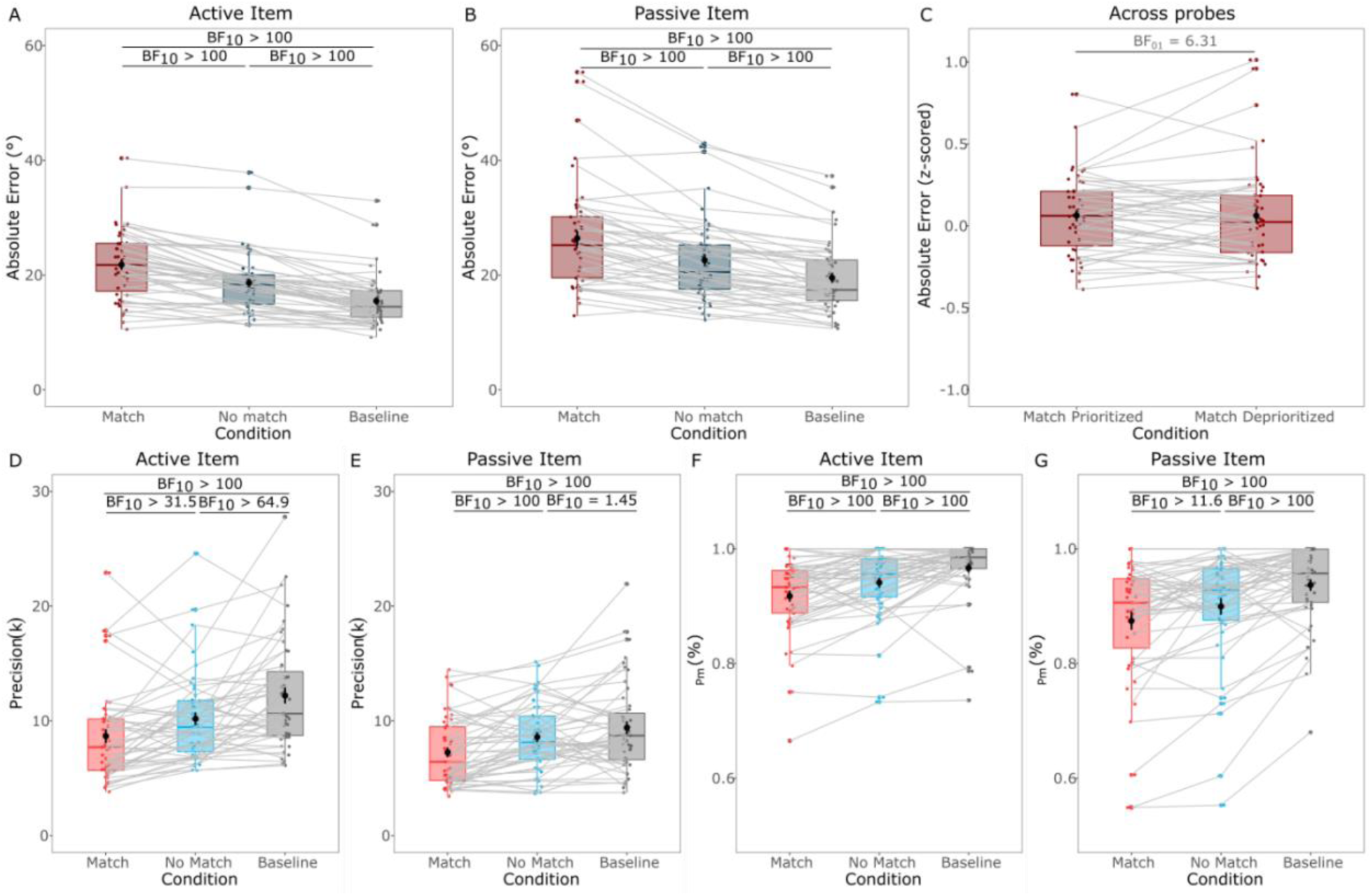
Results for experiment 1. *Note:* A) Absolute error for the active item for the interference versus no interference condition. (B) Absolute error for the passive item for the match versus no match condition. (C) Z-scored absolute error for the match condition across both probes. (D) Precision (*k*) for the active item for match versus no match condition. (E) Precision (*k*) for the passive item for match versus no match condition. (F) Probability of a target response (*P_(m)_*) for the active item for match versus no match condition. (G) Probability of a target response (*P_(m)_*) for the passive item for match versus no match condition. Coloured dots represent individual data points, black dots indicate averages and error bars are +/-SD. Red colour represents the match condition, blue colour represents no match condition. Grey lines connect each subject in the match versus no match condition. Boxplots show median and quartile values.

With respect to the interference, we performed separate repeated-measure Bayesian ANOVAs for the active and passive probes. For the probe of the active item, we found decisive evidence for a main effect of interference (BF_10_ > 100, Figure 2A). We followed this up with one-sided Bayesian t-tests and found decisive evidence for an interference effect for both the match and no match condition relative to the non-interference baseline (*t_paired_* BF_10_ > 100 for both comparisons). We also found decisive evidence that the interference effect was larger in the match compared to the no match condition (*t_paired_* BF_10_ > 100). We performed the same analysis for the passive item where we found decisive evidence for a main effect of interference (BF_10_ > 100, Figure 2B). Using one-sided Bayesian t-tests we again found an interference effect for the match and no match condition, compared to baseline (*t_paired_* BF_10_ > 100 for both comparisons). Furthermore, we found decisive evidence that interference was stronger in the match compared to the no match condition (*t_paired_* BF_10_ > 100). Contrary to our hypothesis, we thus found an interference effect for both the active and the passive item. We therefore wondered if the magnitude of interference differed between the two states. To control for the differences in average error between Probe 1 and Probe 2 we compared the z-scored absolute error between probes and found substantial evidence for the null hypothesis of no difference in the magnitude of interference between states (*t_paired_* BF_10_ = 0.158 (BF_01_ = 6.31), Figure 2C).

As a next step, we were interested in what drives the interference effect, and whether the underlying mechanisms are similar for active and passive items. We therefore fitted two separate standard-mixture models, one for the active item and one for the passive item (Zhang & Luck, 2008). As previously we fitted the model separately for the active and passive item and analysed results using repeated-measures Bayesian ANOVAs, followed by one-sided Bayesian t-tests for post-hoc comparisons.

For the active items we found decisive evidence for a main effect of interference on precision (*k*) (BF_10_ > 100, Figure 2D). Both match and no match conditions showed lower precision compared to the baseline condition with decisive and very strong evidence, respectively (*t_paired_* BF_10_ > 100 and *t_paired_* BF_10_ = 64.95, respectively). Additionally, we found very strong evidence that precision was lower in the match compared to the no match condition (*t_paired_* BF_10_ = 31.5). Furthermore, we found a decisive effect of interference on the probability of target response (*P_(m)_*) (BF_10_ > 100, Figure 2F). Both match and no match conditions showed decisive evidence for an effect on *P_(m)_* relative to the baseline condition (BF_10_ > 100 for both conditions). Additionally, *P_(m)_* was lower in the match condition compared to the no match condition indicated by decisive evidence (BF_10_ > 100).

For the passive item we found decisive evidence for a main effect of interference on precision (BF_10_ > 100, Figure 2E). Both match and no match condition showed evidence of lower precision compared to baseline (*t_paired_* BF_10_ > 100 and *t_paired_* BF_10_ = 1.45, respectively). Notably, the evidence for no match compared to baseline was only anecdotal, but the difference between match and baseline is supported by decisive evidence. In line with this, there was decisive evidence for precision being lower in the match compared to the no match condition (*t_paired_* BF_10_ > 100). We also found a decisive main effect of interference on the probability of a target response (BF_10_ > 100, Figure 2G). The probability of a target response was lower for both the match and the no match condition compared to baseline (*t_paired_* BF_10_ > 100 for both conditions). The probability of a target response was lower in the match compared to the no match condition which is indicated by strong evidence (BF_10_ = 11.62).

Thus, we found that precision, that is, how well an item is represented in memory, and the probability of a target response were similarly affected by the interference manipulation for items that are active or passive within working memory. To further support this interpretation, and to address potential order-of-encoding and cue-validity confounds, we conducted two additional control analyses whose full results appear in the Supplementary Materials. Section S1 (Figure S1) showed that the interference gradient has a similar shape for first- and second-encoded items, with only modest differences in magnitude. Section S2 (Figure S2) demonstrated that the block-wise cue produced a robust, largely additive accuracy benefit across all conditions, which is consistent with the prioritisation manipulation working as intended.

### Experiment 2

In experiment 2 (N=41), we again investigated the effects of interference between active and passive items in working memory; however, this time we interfered in a spatially-specific, rather than feature-specific manner. The results are shown in Figure 3. In accordance with the previous experiment, we found decisive evidence for better performance on the active item (μ_AS_ = 18.9°, sd_AS_ = 5.4°) compared to the passive item (μ_AS_ = 20.9°, sd_AS_ = 4.7°) (*t_paired_* BF_10_ > 100). We also observed decisive evidence that participants performed above chance in the interference task (μ = 95.9%, sd = 7.4%) (*t_one-sided_* BF_10_ > 100).

**Figure 3.**
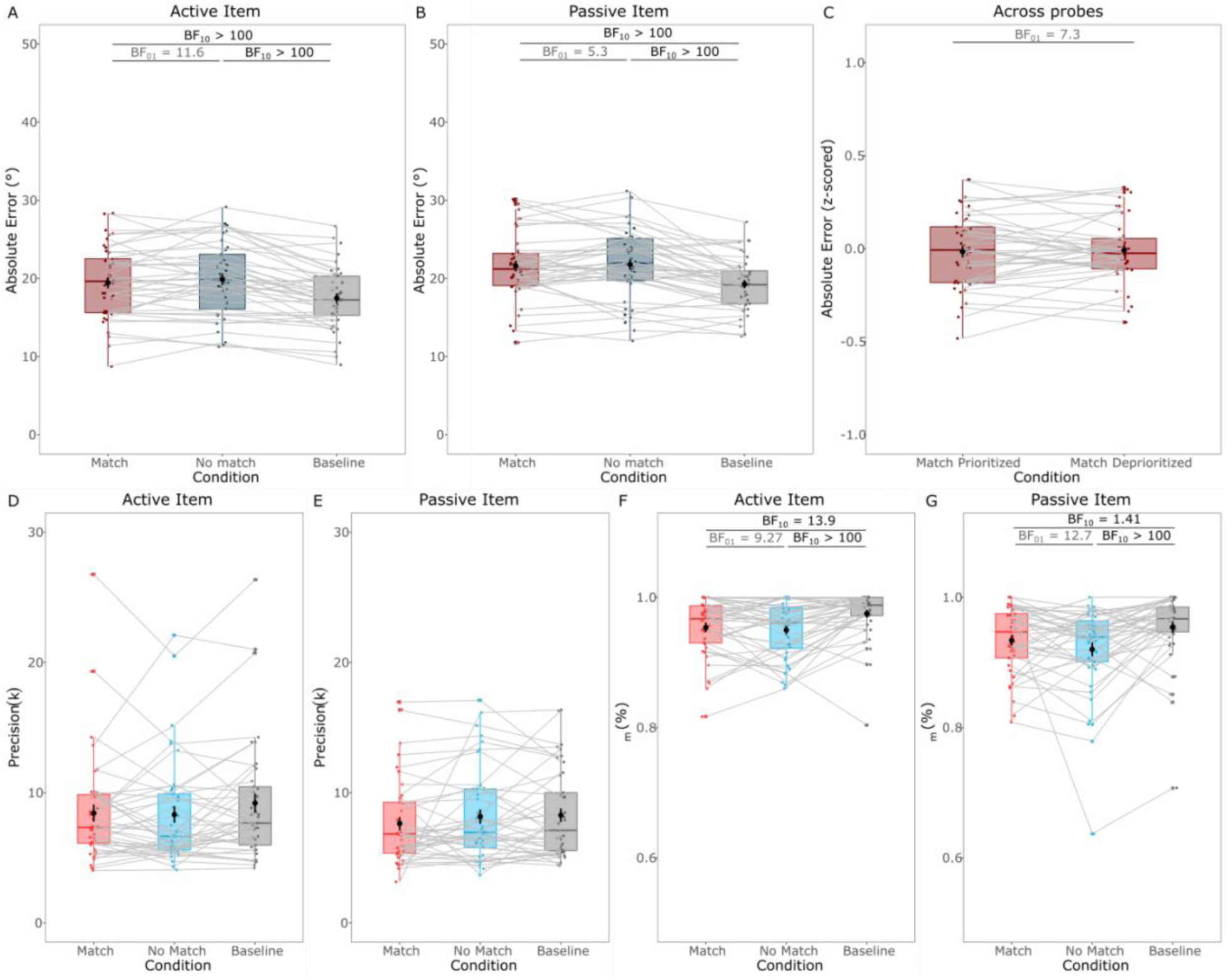
Results for experiment 2. *Note:* (A) Absolute error for the active item for the interference versus no interference condition. (B) Absolute error for the passive item for the match versus no match condition. (C) Z-scored absolute error for the match condition across both probes. (D) Precision (*k*) for the active item for match versus no match condition. (E) Precision (*k*) for the passive item for match versus no match condition. (F) Probability of a target response (*P_(m)_*) for the active item for match versus no match condition (G) Probability of a target response (*P_(m)_*) for the passive item for match versus no match condition. Coloured dots represent individual data points, black dots indicate averages and error bars are +/-SD. Red colour represents the match condition, blue colour represents no match condition. Grey lines connect each subject in the match versus no match condition. Boxplots show median and quartile values.

**Figure 4.**
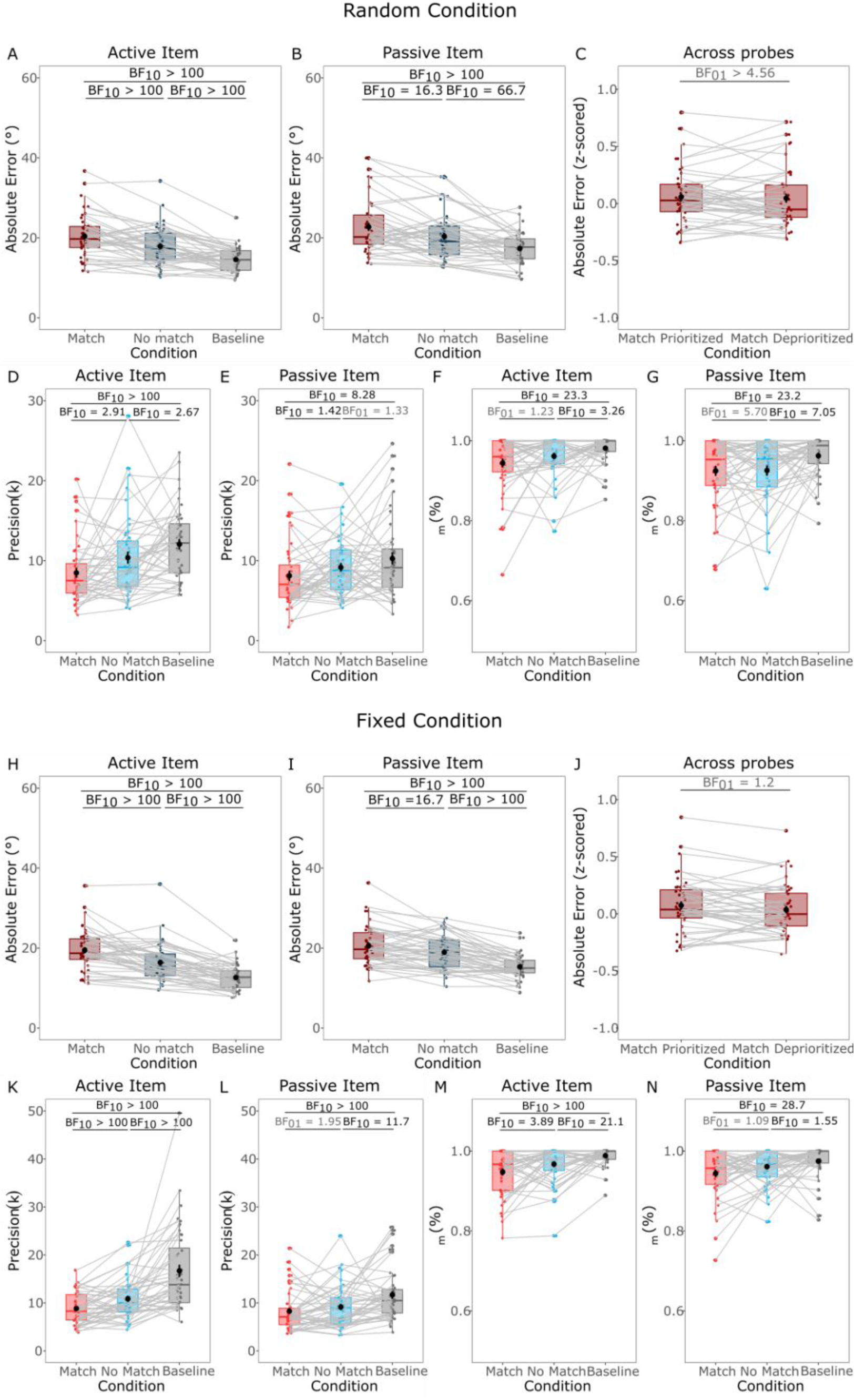
Results of experiment 3. *Note:* Plots **A-G** are for the random condition. (A) Absolute error for the active item for the interference versus no interference condition. (B) Absolute error for the passive item for the match versus no match condition. (C) Z-scored absolute error for the match condition across both probes. (D) Precision (*k*) for the active item for match versus no match condition. (E) Probability of a target response (*P_(m)_*) for the active item for match versus no match condition. (F) Precision (*k*) for the passive item for match versus no match condition. (G) Probability of a target response (*P_(m)_*) for the passive item for match versus no match condition. Plots **H-N** are for the fixed condition. (H) Absolute error for the active item for the interference versus no interference condition. (I) Absolute error for the passive item for the match versus no match condition. (J) Z-scored absolute error for the match condition across both probes. (K) Precision (*k*) for the active item for match versus no match condition. (L) Precision (*k*) for the passive item for match versus no match condition. (M) Probability of a target response (*P_(m)_*) for the active item for match versus no match condition. (N) Probability of a target response (*P_(m)_*) for the passive item for match versus no match condition. Coloured dots represent individual data points, black dots indicate averages and error bars are +/-SD. Red colour represents the match condition, blue colour represents no match condition. Grey lines connect each subject in the match versus no match condition. Boxplots show median and quartile values.

We repeated the same analyses as for experiment 1. We found a decisive main effect of interference on the active item (BF_10_ > 100, Figure 3A). We found decisive evidence of an effect of interference for match and no match condition relative to the baseline (BF_10_ > 100 for both comparisons). However, we found strong evidence against a differential effect between the match and the no match condition (BF_10_ = 0.086 (BF_01_ = 11.64)). Thus, while there was an interference effect relative to the baseline condition; whether the item was presented in the cued location or not did not modulate the interference effect. For the passive item, we also found decisive evidence for a main effect for interference (BF_10_ > 100, Figure 3B). As for the active item, there was again an decisive effect of interference for both the match and no match condition relative to the baseline condition (BF_10_ > 100 for both comparisons). However, we again found substantial evidence against the modulation of the interference effect between the match compared to no match condition (BF_10_ = 0.189; BF_01_ = 5.3). Overall, these analyses suggest that while the interference manipulation worked, because absolute error was higher across both interference conditions when compared to baseline, the interference affected both items to similar extents, independent of spatial location. Comparing the active match condition with the passive match condition further showed that there was substantial evidence that there is no difference in the amount of interference between them (BF_10_ = 0.137 (BF_01_ = 7.3, Figure 3C).

As a next step, we looked at the model parameters of the standard-mixture model. We found only anecdotal evidence that there was no main effect on precision for the active item (BF_10_ = 0.6 (BF_01_ = 1.658), Figure 3D). We did find evidence for a very strong main effect on the probability of a target response for the active item (BF_10_ = 79.42, Figure 3F). The probability of a target response was lower for the match and no match condition compared to baseline supported by strong and decisive evidence (BF_10_ = 13.97 and BF_10_ > 100, respectively). There was substantial evidence for no effect on the probability of a target response between the match and no match condition (BF_10_ = 0.11 (BF_01_ = 9.27)). For the passive item there was again substantial evidence for no effect on precision either (BF_10_ = 0.21 (BF_01_ = 4.82), Figure 3E). There was substantial evidence for a main effect on the probability of a target response for the passive item (BF_10_ = 6.9, Figure 3G). We found only anecdotal evidence for a lower probability of a target response for the match condition compared to baseline (BF_10_ = 1.41), but we did find decisive evidence for a difference between the no match condition and the baseline (and BF_10_ > 100), and strong evidence for a lack of effect for a difference between match and no match condition BF_10_ = 0.079 (BF_01_ = 12.7)). Overall, we did not find any evidence in this study for location-specific modulation of interference, and neither did we find evidence for a modulation of interference between the functional states of the items in working memory. The general interference effect between both interference conditions compared to the baseline that we did find was driven by a decrease in the probability of a target response, which means there was increased guessing of the response by participants.

### Experiment 3

In experiment 3 (N=44), we wanted to test if expectations about the upcoming interference could modulate the effect. Since we had a stronger interference effect in experiment 1 compared to experiment 2, we went back to feature-based interference. This time we did not only block the item priority, but we also included a blocked and explicit cue for the interference feature. Thus, participants could anticipate which item was going to be interfered with and we kept this constant across each block of trials. The idea was that this would provide an opportunity to the participants to exert strategic control over memory maintenance, if they could, possibly depending on the functional state of the items being maintained.

We found decisive evidence for higher accuracy for the active item (μ_AS_ = 16.9°, sd_AS_ = 5.5°) compared to the passive item (μ_AS_ = 19.2°, sd_AS_ = 5.3°) (*t_paired_* BF_10_ > 100). As in previous experiments, participants accurately responded in the interference condition supported by decisive evidence, indicating that they paid attention to the task (μ = 89.9%, sd = 7%) (*t_one-sided_* BF_10_ > 100).

Again, we performed the same analysis as in the previous experiment, but we separated the blocks into random interference blocks (identical to experiment 1) and fixed interference blocks, where the interference condition stayed the same throughout the experiment and was cued to the participant.

#### Random condition

We found a decisive main effect of interference on the active item (BF_10_ > 100). We found decisive evidence for an effect of interference on both the match and no match condition compared to baseline (BF_10_ > 100 for both comparisons). Furthermore, we found decisive evidence for an effect of interference between the match and no match condition (BF_10_ > 100). We found the same decisive main effect of interference for the passive item (BF_10_ > 100). Interference was larger for both the match and no match condition relative to the baseline condition supported by decisive and very strong evidence (BF_10_ > 100 and BF_10_ = 66.75, respectively). Additionally, strong evidence indicates that interference was larger for the match compared to the no match condition (BF_10_ = 16.3). As before, we found substantial evidence that the interference effect was not modulated by the functional state of the items in working memory (BF_10_ = 0.22 (BF_01_ = 4.56)).

Fitting the standard-mixture model to this data, we found a decisive main effect of interference on precision for the active item (BF_10_ > 100). Precision was notably lower in the match condition compared to baseline, which was supported by decisive evidence (BF_10_ > 100); however, the effect of the no match condition relative to baseline was more unclear as we found only anecdotal evidence for the effect (BF_10_ = 2.67). Furthermore, anecdotal evidence shows that precision was potentially lower in the match compared to the no match condition (BF_10_ = 2.91). There was also a strong main effect of interference on the probability of a target response for the active item (BF_10_ = 10.12). We found strong evidence that the match condition had a lower probability of a target response compared to baseline (BF_10_ = 23.31) but again observed only anecdotal evidence for a difference between the no match condition relative to baseline (and BF_10_ = 3.26). Unlike the clear evidence for a difference between match and no match condition in experiment 1, in this experiment the evidence in favour or against a difference between the match and no match condition was only anecdotal (BF_10_ = 0.757 (BF_01_ = 1.32)). For the passive item, we found anecdotal evidence for a main effect of interference on precision (BF_10_ = 2.84). Post-hoc comparisons did show substantial evidence that the match condition had lower precision compared to the baseline (BF_10_ = 8.28); however, we found anecdotal evidence that there was no difference between the no match condition and the baseline (BF_10_ = 0.75 (BF_01_ = 1.33)), and also just anecdotal evidence for a difference between the match and no match condition on precision (BF_10_ = 1.42). Overall, the effect of precision on the passive item is largely found by anecdotal evidence and therefore inconclusive. There was substantial evidence for a main effect on the probability of a target response (BF_10_ = 5.26). We found strong and substantial evidence for reduced probability of a target response in the match and no match condition relative to baseline (BF_10_ = 23.32 and BF_10_ = 7.05, respectively). However, we found substantial evidence against a difference between the match and no match condition on the probability of a target response (BF_10_ = 0.173 (BF_01_ = 5.79)).

Overall, we observed some discrepancies in the model parameters when compared to experiment 1. However, these were only substantiated by anecdotal evidence. Mainly, we found anecdotal evidence for the null hypothesis in two cases where we previously found strong evidence for the alternative hypothesis in experiment 1. These findings could be caused by the reduced trial numbers of this part of the experiment compared to experiment 1, which was due to the second condition that we included here – especially since the evidence for the null hypothesis was only anecdotal. Notably we did find substantial evidence for no difference between the match and no match condition on the probability of a target response for the passive item, which was different from the results found in experiment 1.

#### Fixed condition

There was a decisive main effect of interference for the active item (BF_10_ > 100). We found decisive evidence for a larger error in the match and no match interference condition compared to baseline (BF_10_ > 100 for both comparisons), as well as decisive evidence for a larger error in the match compared to the no match condition (BF_10_ > 100). For the passive item we found the same decisive main effect of interference (BF_10_ > 100). Again, the error was larger in the match and no match condition compared to baseline supported by decisive evidence (BF_10_ > 100 for both comparisons), and strong evidence showed a larger error in the match compared to the no match condition (BF_10_ = 16.68). We found anecdotal evidence that the interference effect was not modulated by the priority state in working memory (BF_10_ = 0.93 (BF_01_ = 1.2)), leaving open the possibility that with more power in the fixed-condition version of this task the magnitude of the interference effect might change depending on priority state.

We found a decisive main effect of interference on precision for the active item (BF_10_ > 100). Decisive evidence supports that precision was lower for both match and no match interference conditions compared to baseline (BF_10_ > 100 for both comparisons). Decisive evidence also shows that precision was lower for the match compared to the no match condition (BF_10_ > 100). Interference also showed decisive evidence for a main effect on the probability of a target response (BF_10_ > 100). We found decisive and strong evidence for a lower probability of a target response in both the match and no match condition compared to the baseline condition (BF_10_ > 100 and BF_10_ = 21.13, respectively). We also found substantial evidence for lower probability of a target response in the match compared to the no match condition (BF_10_ = 3.89). For the passive item, we found a decisive main effect of interference on precision (BF_10_ > 100). We found evidence of lower precision for the match and no match condition relative to baseline supported by decisive and strong evidence (BF_10_ > 100 and BF_10_ = 11.75, respectively). We found some evidence against a difference between the match and no match condition on precision, but this was only supported by anecdotal evidence (BF_10_ = 0.514 (BF_01_ = 1.95). We also found substantial evidence of a main effect on the probability of a target response (BF_10_ = 8.6). We found strong evidence that the probability of a target response was lower in the match compared to the baseline condition (BF_10_ = 28.77), as well as anecdotal evidence for a difference between the match and no match condition (BF_10_ = 1.55). We found anecdotal evidence against a difference between the no match and baseline condition (BF_10_ = 0.92 (BF_01_ = 1.09)).

Overall, the patterns we observed were very similar between the fixed and random condition trials, indicating that participants did not benefit from having explicit knowledge of the interference condition, and that active and passive items were still equally affected by interference feature-based interference. Following this observation, we finally tested if participants had any benefit from knowledge of the interference feature. To this end, we calculated the difference between interference (match and no match condition combined) minus the baseline condition to test if the magnitude of interference was modulated by the fixed versus random condition. We then performed a paired-sample t-test between these difference scores for the active and passive item separately. We found anecdotal evidence that the magnitude of interference did not differ between random and fixed blocks for the active item (BF_10_ = 0.36 (BF_01_ = 2.75)) and substantial evidence that the magnitude did not differ for the passive item (BF_10_ = 0.17 (BF_01_ = 5.87)). Thus, while we can be sure that the passive item did not show a differential effect between random and fixed blocks, it remained inconclusive if this was also the case for the active item.

### Experiment 4

In experiment 4 (N=56), we changed the nature of the block cue; rather than cueing the response order to achieve a prioritised and deprioritised item, we now associated the cued item with higher monetary reward for correct performance compared to the uncued item. This allowed us to achieve prioritised by reward association and randomised response order. As previously we found definitive evidence for higher accuracy for the item responded to first (Probe 1), compared to the item responded to second (Probe 2) (μ_AS_ = 16.2°, sd_AS_ = 3.39° and μ_AS_ = 17.6°, sd_AS_ = 3.75°, respectively) (*t_paired_* BF_10_ > 100). This experiment no longer potentially confounds response order with prioritisation, so we also compared response accuracy for the active item (prioritised; higher rewarded) with the passive item (deprioritised; lower rewarded), and found substantial evidence for higher accuracy for the active item (μ_AS_ = 16.5°, sd_AS_ = 3.25°), compared to the passive item (μ_AS_ = 17.3°, sd_AS_ = 3.87°) (*t_paired_* BF_10_ = 6.09). Finally, to ensure that cue status and response order did not interact with each other in this design, we performed a 2 (Cue status) x 2 (Response order) Bayesian repeated measures ANOVA. We found that the best model included Cue status and Probe order as a main effect (BF_10_ > 100; see Supplementary Table 13 for the full output) for this model with decisive evidence for Probe order (BF_incl_ > 100) and substantial evidence for Cue status (BF_incl_ = 5.16) that these factors contribute to the model. We found no evidence that an interaction effect would further contribute to the model (BF_incl_ = 0.65). Thus, together we interpret this result as indicating that the cued item was indeed in a prioritised state leading to small, but statistically robust higher performance. We further observed decisive evidence that participants performed above chance in the interference task (μ = 89.4%, sd = 4.2%) (*t_one-sided_* BF_10_ > 100), indicating that participants paid attention to the interference stimuli.

As previously, we next wanted to investigate the effect of interference on the active and passive items. Using the same repeated-measure Bayesian ANOVA approach as before, we found decisive evidence for a main effect of interference on the active item (BF_10_ > 100, Figure 5A). We followed this up with one-sided Bayesian t-tests and found decisive evidence for an effect of interference for both the match and no match condition, relative to the baseline condition (BF_10_ > 100 for both comparisons). We further found decisive evidence for larger interference in the match compared to the no match condition (BF_10_ > 100). Doing the same analyses for the passive item we again found definitive evidence for a main effect of interference (BF_10_ > 100, Figure 5B). Post-hoc tests showed definitive evidence for interference in the match and no match condition relative to baseline (BF_10_ > 100 for both comparisons), as well as more interference for the match compared to the no match condition (BF_10_ > 100). Finally, since we found interference for both items we also tested if the magnitude of the interference effect differed between the states. Using z-scored absolute errors to account for accuracy differences between the active and passive item we found anecdotal evidence that there was no difference in the magnitude of the interference effect (*t_paired_* BF_10_ = 0.211 (BF_01_ = 4.75), Figure 5C). Thus, also after changing our design to detangle response order from priority state, and thereby ensuring that our block manipulation led to a prioritisation of items within WM, we still found no effect of protection from interference for passive – presumably silent – items within WM, in line with our results of the previous three experiments.

**Figure 5.**
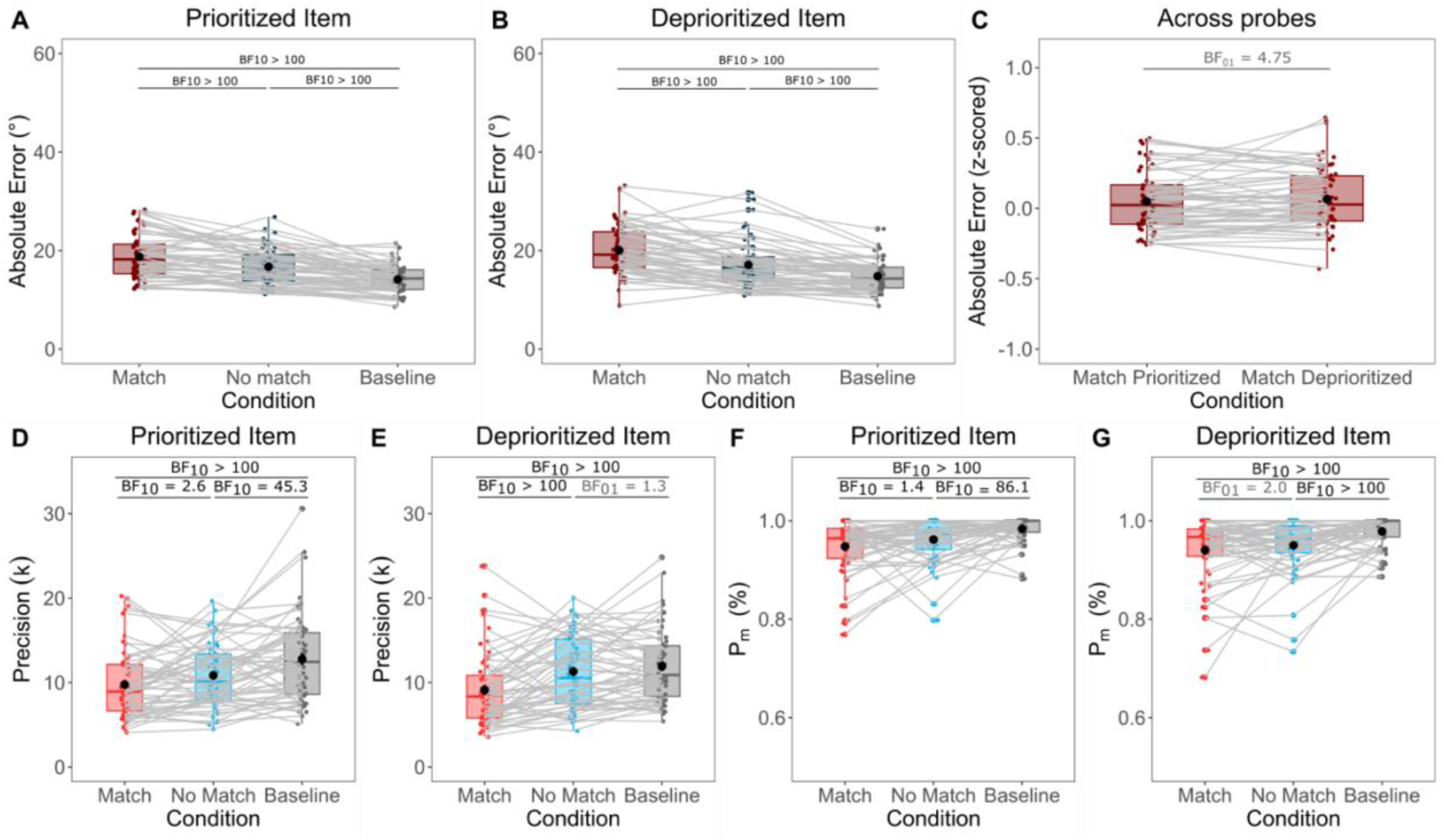
Results of experiment 4. *Note:* (A) Absolute error for the active item for the interference versus no interference condition. (B) Absolute error for the passive item for the match versus no match condition. (C) Z-scored absolute error for the match condition across both probes. (D) Precision (*k*) for the active item for match versus no match condition. (E) Precision (*k*) for the passive item for match versus no match condition. (F) Probability of a target response (*P_(m)_*) for the active item for match versus no match condition (G) Probability of a target response (*P_(m)_*) for the passive item for match versus no match condition. Coloured dots represent individual data points, black dots indicate averages and error bars are +/-SD. Red colour represents the match condition, blue colour represents no match condition. Grey lines connect each subject in the match versus no match condition. Boxplots show median and quartile values.

To ensure this also holds when looking at the results in more details, we went on to fi t the standard-mixture model as we did for previous experiments. For the active item we found a main effect of interference on precision (*k*) (BF_10_ > 100, Figure 5D). We found at least very strong evidence that the match and no match condition had lower precision compared to the baseline (BF_10_ > 100 and BF_10_ = 45.3, respectively). However, evidence was only anecdotal for a difference between the match and no match condition (BF_10_ = 2.6). For the probability of a target response (*P_(m)_*) we found decisive evidence for an effect of interference condition (BF_10_ > 100, Figure 5F). Post-hoc tests revealed at least very strong evidence for an effect of guess rate on the match and no match condition when compared to the baseline condition (BF_10_ > 100 and BF_10_ = 86.1, respectively). However, again there was no clear evidence for a differential effect of guess rate between the match and no match condition (BF_10_ = 1.4).

For the passive item, we repeated the same analyses as above. We found decisive evidence for a main effect of interference on precision (BF_10_ > 100, Figure 5E). We found decisive evidence that precision was lower for the match condition compared to baseline (BF_10_ > 100); however, we found no clear evidence for a difference between baseline and the no match condition in either direction (BF_10_ = 0.76 (B_01_ = 1.3)). The match condition did show lower precision compared to the no match condition, which was supported by decisive evidence (BF_10_ > 100). For guess rate we found decisive evidence for an effect of interference (BF_10_ > 100, Figure 5G). We also found decisive evidence for a difference between match and no match condition compared to baseline (BF_10_ > 100 for both conditions). However, we found no clear evidence for a difference between match and no match condition on guess rate (BF_10_ = 0.5 (BF_01_ = 2.0)). Thus, we show that overall, the pattern remains as shown in previous experiments, even when cueing priority in a different manner that excluded the possibility of a confound of response order and cue status.

## Discussion

Working memory can be stored in persistent-activity or activity-quiescent states, and an accumulating amount of evidence has shown that these two mechanisms of WM storage are not mutually exclusive (Lundqvist, Herman, & Miller, 2018; Muhle-Karbe et al., 2021; Oberauer & Awh, 2022). However, little research to date has focused on the potential functional roles of these differing states in WM. In the current study, we set out to test one potential functional role of these varying states – namely, the hypothesis that functionally passive (and neuronally activity-quiescent) states could be protecting items from interference, compared to functionally active, (and neuronally persistent-activity) states.

For this, we tested the effect of an interference task presented during WM maintenance in three experiments. We were primarily interested in potential differences between the maintenance of active (item cued to be probed first and therefore thought to be prioritised) and passive (item cued to be probed second and therefore thought to be deprioritised) items, as they might be supported by different neural mechanisms. Active items might be held by means of persistent neural firing, which could be mediated by internal attention (Bae & Luck, 2018; Lewis-Peacock & Postle, 2012). Conversely, passive items might be retained in an activity-quiescent manner, through functional connectivity that results from short-term synaptic plasticity (Mongillo et al., 2008; Stokes, 2015). We hypothesised that activity-quiescent maintenance through functional connectivity might afford better protection from interference (Kozachkov et al., 2022; Lundqvist, Herman, Warden, et al., 2018; Miller et al., 2018).

Contrary to our hypothesis, we found no evidence that passive items (assumed to be maintained in a quiescent state) were more protected from interference compared to active items (assumed to be in a persistent-activity state). Instead, both states showed similar susceptibility to interference across all three experiments, regardless of whether the distractor overlapped in feature (Experiment 1) or spatial dimension (Experiment 2), a pattern that persisted even when the participants had explicit knowledge about the nature of the upcoming interference (Experiment 3) or when we changed the cueing paradigm to a reward based regime (Experiment 4). Our findings are in line with recent results from Zhang and Lewis-Peacock (2023), who similarly found no difference in interference susceptibility between active and passive items – using face stimuli varying along sex and age dimensions. Additionally, this is further in line with recent findings by Hautekiet and colleagues (2024) who investigated prioritisation modes using colour-shape sequences and reward-based prioritisation to show that prioritised items are not more shielded from distractors.

At the risk of pointing out the obvious, our data clearly also do not support the idea that prioritised items might *heighten* susceptibility to certain forms of disruption (Hitch et al., 2018; Hu et al., 2014; Mallett & Lewis-Peacock, 2019), nor that prioritised items always enjoy protection from interference (Makovski et al., 2008; Makovski & Pertzov, 2015; Schneider et al., 2017; Souza & Oberauer, 2016). Indeed, a recent study has demonstrated that different prioritisation manipulations – test-relevance cuing, reward incentives and refreshing – yield distinct interference profiles, highlighting that not all priority signals engage the same protective mechanisms (Vergauwe et al., 2025).

What could be causing such seemingly contradictory outcomes regarding the nature of interference effects in previous studies? One possibility involves the specifics of the task – for instance, the exact nature of the distractor (Hu et al., 2014; Pinto et al., 2013), the number of to-be-remembered items (Allen & Ueno, 2018), or the time points of distractor presentation. All of these could determine whether priority confers resilience or risk. Additionally, how “priority” is operationalised – through retro-cued, reward or attentional modulations – may lead to divergent results. Explicit perceptual-comparison judgments amplify memory biases induced by new inputs, while at the same time prioritisation has been shown to retroactively reduce distortion, yet enhances proactive perceptual biases (J. Saito et al., 2025; J. M. Saito et al., 2023). Such bidirectional interactions might modulate how interference tasks impact WM representations. Here, we used explicit blocks to designate which item was relevant first versus second, offering a relatively direct manipulation of priority. This approach aligns with Muhle-Karbe and collagues (2021), who showed that prioritised items display heightened neuronal representations while deprioritised ones can exist in a silent trace. Nevertheless, in our case, the state the items were in did not lead to a differential effect of interference.

In addition to the question of whether deprioritised items are more protected from interference, our data also allow for a test of whether interference might be different between spatial and featural overlap. Experiment 1 (feature-based interference) showed a robust match effect on accuracy and precision: Both active and passive items showed higher disruption when the distractor shared the same colour or orientation. Interestingly, in Experiment 2 (space-based interference), the match versus no-match distinction was less critical. The presence of a spatial distractor affected WM performance overall, when compared to the no distractor baseline condition, but matching locations did not systematically enhance interference. This outcome aligns with domain-specific interference theories, whereby spatial overlap may be less disruptive if the stored features (colour/orientation) do not critically rely on spatial codes (Awh & Jonides, 2001; Jerde et al., 2012).

Regardless of the specific dimension, the critical finding remains that the magnitude of interference did not systematically differ between active and passive states. This was further extended in Experiment 3, where participants had explicit knowledge about which feature would be targeted by the interference task. Even when provided with this opportunity to anticipate the nature of the interference task, our data suggests that participants were not able to make use of this strategic cue to protect the working memory content in either state from interference. Thus, while interference can take place in multiple forms – feature or space-based – and can be anticipated, none of these manipulations selectively provided increased resilience of passive items. These findings are in line with a recent study that similarly reported the absence of robust protection of quiescent states in working memory (Hautekiet et al., 2025).

If these presumably quiescent states of passive items in WM do not afford robust protection from external distractors, why might the brain maintain items in silent traces? One possibility could be that quiescent states help minimise internal interference among multiple items stored in WM (Chatham & Badre, 2015; Rose et al., 2016). By storing passive items – which are only relevant later in time – in a synaptic format, the system might free up active firing for the currently most relevant item, thereby reducing crosstalk (Muhle-Karbe et al., 2021). Indeed, it is a well-accepted fact that WM capacity is strongly limited, which would suggest that it is important to dynamically balance activity across items, and having only one, or few items fully “activated” at a time. This mechanism may be especially crucial under higher loads than tested here (only two items). Future work could explore whether the functional benefit of quiescent states is relevant for preventing overlap within WM itself, rather than insulating those items against external interference.

The current study comes with some limitations that should be noted. First, we did not take any direct neural measures, which means that our interpretations as to potential underlying mechanisms remain indirect inferences based on previous papers (e.g., LaRocque et al., 2012; Muhle-Karbe et al., 2021; Wolff et al., 2017). Specifically, we cannot reliably determine whether our prioritisation manipulation successfully led to prioritised items being stored in a neuronally distinct and more active format compared to passive items. Future research should incorporate neural measures to directly investigate how WM content is represented under different cue states within this paradigm. This is particularly important given accumulating evidence that passive representations may not protect against interference. However, direct neural evidence for this effect is still lacking, leaving open the possibility that prioritisation manipulations have generally been unsuccessful across experiments (Hautekiet et al., 2025; Z. Zhang & Lewis-Peacock, 2023). In relation to this, with purely behavioural measures we cannot unambiguously determine which item occupied the focus of attention during maintenance. The small quantitative differences between first- and second-encoded items (Supplementary analyses) therefore leave open the possibility that recency or focus-of-attention effects contributed to the observed interference pattern, rather than a “pure” difference between active and passive states. Although Experiment 4 showed that the cue manipulation produces a reliable priority benefit independent of probe order, future work combining this paradigm with neural measures will be needed to track attentional focus directly and test how it relates to interference. Second, we only tested memory for a pair of items, which is a relatively modest load on working memory, and which might not have required offloading of information in a quiescent state. A higher WM load might reveal different patterns: If load increases, there might be more need for items to be offloaded in a quiescent state to avoid cross-item interference. Third, although we manipulated whether participants were cued to the nature of the interference task (Experiment 3), the precise temporal dynamics of the interference may play a major role in whether silent states become reactivated or remain quiescent, which we did not modulate. Future paradigms might systematically vary timing, load, and distractor complexity to better clarify when—if ever—silent states reduce interference from external inputs. Finally, it should be noted that overall performance on our task was rather high, which could have a potential ceiling effect on accuracy. While we do see a consistent effect of interference relative to the baseline condition – suggesting that our manipulation was still strong enough to decrease accuracy in response to interference – we cannot exclude that results might differ in a more difficult task setting.

To conclude, across three experiments involving feature-based and space-based interference, we did not see protection from interference on passive items in WM. Even explicit knowledge of which feature dimension would be targeted by the interference task did not protect passive (nor active) items from interference. Rather than serving as an external interference shield, quiescent states may be functionally critical for mitigating internal interference among concurrently stored items. Future work that combines higher WM loads, neural measures, and different interference paradigms could further elucidate the possible functional advantages of short-term synaptic plasticity in WM, and clarify how the brain flexibly transitions items between active and quiescent states to meet external demands.

## Author contributions

Sophia Wilhelm contributed to conceptualisation of the experiments, and served as lead for data curation, formal analysis, methodology, visualisation and writing – original draft. Yuanyuan Weng contributed to data curation and reviewed the final manuscript. Jelmer Borst and Robbert Havekes reviewed the final manuscript. Elkan Akyürek served as lead to conceptualisation of the experiments, project administration and supervision and contributed to writing – original draft.

