## Supplementary Materials for "Feature and space-based interference with functionally active and passive items in working memory"

### 1. Presentation order modulation of interference (Experiment 1-3)

To determine whether the magnitude of the distractor-induced interference depends on the time since an item has been encoded, we re-analysed the data as a function of presentation order. We did this based on data from Experiment 1 which shows the strongest interference effect, for data from Experiment 2 which has a slightly changed design which a distinct spatial location for each encoded item, and finally for Experiment 3.

**Experiment 1:**

We calculated a 2 (presentation order: first-encoded versus second-encoded item) x 3 (interference condition: baseline, no-match, match) repeated-measures ANOVA. This analysis yielded a reliable interaction between presentation order and interference condition ($F$ (1.76, 82.54) = 4.78, $p$ = 0.015; $BF_{10}$ > 100) (For full results see Table 1+2). However, as evidenced by Figure S1A, the mean absolute error increased monotonically from the baseline condition to the no-match condition and further to the match condition. Nevertheless, it should be noted that that the size of this stepwise increase was modestly greater when the item had been encoded second, indicating that distractor interference is amplified when the memorised feature is more recently encoded.

***Supplementary Table 1***

Full ANOVA table for the frequentist analysis of Experiment 1.

| Effect | DF | MSE | F | p-value |
|---|---|---|---|---|
| Presentation order | 1,47 | 17.56 | 8.36 | 0.006 |
| Interference condition | 1.48, 69.61 | 12.77 | 103.82 | <0.001 |
| Presentation order * interference condition | 1.76, 82.54 | 6.36 | 4.73 | 0.015 |

***Supplementary Table 2***

Full ANOVA table of Bayesian analysis of Experiment 1. Table ordered from best to worst model.

| Model | P(M) | BF(M\|data) | $BF_M$ | $BF_{10}$ | Error (%) |
|---|---|---|---|---|---|
| Null model (incl. subject + random slope | 0.2 | $1.075x10^{-23}$ | $4.300x10^{-23}$ | 1 | |
| Presentation order + Interference condition + Presentation Order * Interference condition | 0.2 | 0.712 | 9.911 | > 100 | 4.51 |
| Presentation order + Interference condition | 0.2 | 0.261 | 1.415 | > 100 | 23.12 |
| Interference condition | 0.2 | 0.026 | 0.108 | > 100 | 3.783 |
| Presentation order | 0.2 | $7.454x10^{-23}$ | $2.982x10^{-22}$ | 6.933 | 4.797 |

*Analysis of Effects (across all models)*

| Effects | P(incl) | P(excl) | P(incl\|data) | P(excl\|data) | $BF_{incl}$ |
|---|---|---|---|---|---|
| Presentation order | 0.6 | 0.4 | 0.974 | 0.026 | 24.762 |
| Interference condition | 0.6 | 0.4 | 1 | $3.442x10^{-15}$ | $1.937x10^{+14}$ |

| | | | | | |
|---|---|---|---|---|---|
| Presentation order * Interference condition | 0.2 | 0.8 | 0.712 | 0.288 | 9.911 |

**Experiment 2:**

We repeated the same analysis as we did for Experiment 1 above on the data of Experiment 2. In Experiment 2 the items were presented in a unique spatial location instead of centralised, which might change the encoding dynamics, and thus we wanted to replicate the same analysis on this data. Indeed, we only found an effect of interference condition ($F$ (1.69, 67.70) = 32.87, $p$ < 0.001; $BF_{10}$ > 100; Figure S1B), but no main effect for presentation order or an interaction between interference condition and presentation order as we had observed in Experiment 1 (See table 3 for frequentist statistics and table 4 for full Bayesian models). Thus, in this experiment the more recently encoded item was not more vulnerable to interference.

***Supplementary Table 3***

Full ANOVA table for the frequentist analysis of Experiment 2

| **Effect** | **DF** | **MSE** | **F** | **p-value** |
|---|---|---|---|---|
| Presentation order | 1.40 | 13.01 | 2.62 | 0.113 |
| Interference condition | 1.69,67.70 | 5.27 | 32.87 | <0.001 |
| Presentation order * interference condition | 1.97,78.83 | 3.71 | 0.7 | 0.496 |

***Supplementary Table 4***

Full ANOVA table of Bayesian analysis of Experiment 2. Table ordered from best to worst model.

| **Model** | **P(M)** | **BF(M\|data)** | **$BF_M$** | **$BF_{10}$** | **Error (%)** |
|---|---|---|---|---|---|
| Null model (incl. subject + random slope | 0.2 | 5.811×10$^{-9}$ | 2.324×10$^{-8}$ | 1.0 | |
| Interference condition | 0.2 | 0.526 | 4.436 | 9.050×10$^{+7}$ | 1.596 |
| Presentation order + Interference condition | 0.2 | 0.415 | 2.832 | 7.134×10$^{+7}$ | 2.432 |
| Presentation order + Interference condition + Presentation order * Interference condition | 0.2 | 0.060 | 0.254 | 1.027×10$^{+7}$ | 2.292 |
| Presentation order | 0.2 | 4.276×10$^{-9}$ | 1.711×10$^{-8}$ | 0.736 | 2.174 |

*Analysis of Effects (across all models)*

| **Effects** | **P(incl)** | **P(excl)** | **P(incl\|data)** | **P(excl\|data)** | **$BF_{incl}$** |
|---|---|---|---|---|---|
| Presentation order | 0.6 | 0.4 | 0.47 | 0.526 | 0.601 |
| Interference condition | 0.6 | 0.4 | 1 | 1.009x10$^{-8}$ | 6.609x10$^{+7}$ |
| Presentation order * Interference condition | 0.2 | 0.8 | 0.06 | 0.94 | 0.254 |

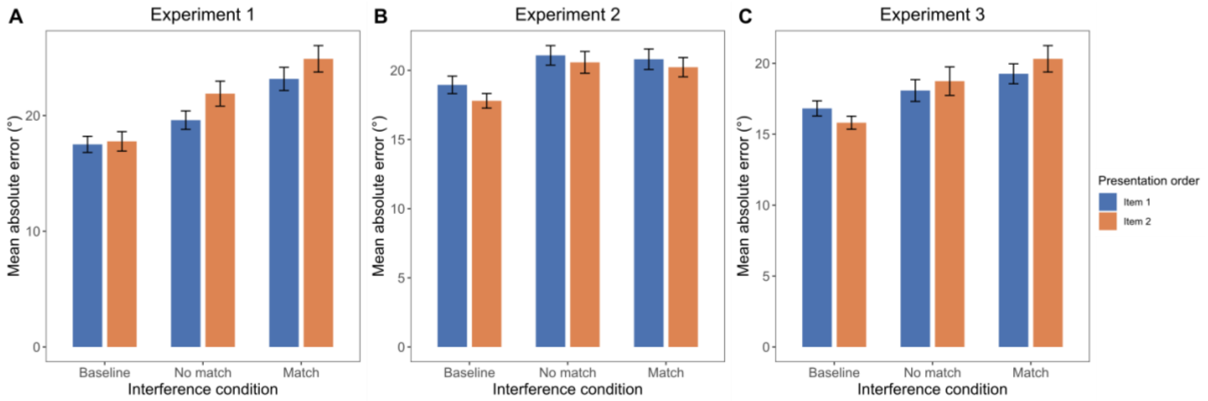


*Figure S1:* Absolute error across the three interference conditions in Experiment 1 (**A**), Experiment 2 (**B**) and Experiment 3 (**C**) (baseline, no match and match), depending on if the item was presented first or second – independent of the prioritisation cue. Boxplot shows the mean, and error bars indicate the SEM.

**Experiment 3:**

Finally, we repeated the same analysis as we did for the two previous experiments also for Experiment 3. As for Experiment 1 we found a significant interaction between presentation order and interference condition ($F$ (1.99, 85.67) = 4.52, $p$ = 0.014; Figure S1C). However, using Bayesian model comparisons we found that this interaction effect only provided the second-best fitting model, while the best fitting model only contained interference condition ($BF_{10} > 100$) with only anecdotal evidence for an additional effect of the interaction term ($BF_{incl}$ = 2.183) (See table 5 for frequentist statistics and table 6 for full Bayesian models). Thus, while we consistently found an effect of interference condition on absolute error; the data of this experiment did not provide us with definite evidence for an interaction of presentation order and interference condition.

***Supplementary Table 5***

Full ANOVA table for the frequentist analysis of Experiment 3.

| **Effect** | **DF** | **MSE** | **F** | **p-value** |
|---|---|---|---|---|
| Presentation order | 1,43 | 12.49 | 0.29 | 0.592 |
| Interference condition | 1.64,70.61 | 27.22 | 12.05 | <0.001 |
| Presentation order * interference condition | 1.99,85.67 | 5.86 | 4.52 | 0.014 |

***Supplementary Table 6***

Full ANOVA table of Bayesian analysis of Experiment 3. Table ordered from best to worst model.

| **Model** | **P(M)** | **BF(M\|data)** | **$BF_M$** | **$BF_{10}$** | **Error (%)** |
|---|---|---|---|---|---|
| Null model (incl. subject + random slope | 0.2 | $5.384 \times 10^{-4}$ | 0.002 | 1.000 | |
| Interference condition | 0.2 | 0.528 | 4.481 | 981.342 | 2.036 |
| Presentation order + Interference condition + Presentation Order * Interference condition | 0.2 | 0.353 | 2.183 | 655.806 | 6.629 |
| Presentation order + Interference condition | 0.2 | 0.118 | 0.534 | 218.896 | 4.872 |

| | | | | | |
|---|---|---|---|---|---|
| Presentation order | 0.2 | $1.213\times10^{-4}$ | $4.851\times10^{-4}$ | 0.225 | 3.906 |

*Analysis of Effects (across all models)*

| **Effects** | **P(incl)** | **P(excl)** | **P(incl\|data)** | **P(excl\|data)** | **$BF_{incl}$** |
|---|---|---|---|---|---|
| Presentation order | 0.6 | 0.4 | 0.471 | 0.529 | 0.594 |
| Interference condition | 0.6 | 0.4 | 0.999 | $6.597\text{x}10^{-4}$ | 1009.904 |
| Presentation order * Interference condition | 0.2 | 0.8 | 0.353 | 0.647 | 2.183 |

Together, these analyses showed that across both experiments, the first-encoded and second-encoded items were both susceptible to distractor interference. In Experiment 1 and 3 there was some evidence that this interference is modestly amplified for more recently encoded items, whereas this modulation by presentation order did not emerge in Experiment 2. Thus, while we acknowledge that there is a potential effect of order, these analyses do not alter our central conclusions.

**2. Effectiveness of the block-wise prioritisation cue (Experiment 1-3)**

To verify that the block-wise cue manipulation successfully prioritised the to-be-probed item – beyond any advantage conferred by temporal proximity to the probe – we re-analysed Experiment 1, 2 and 3 as a function of cue status, for item 1 and item 2 separately. We performed a 2 (cue status: cued versus uncued) x 2 (presentation order: item 1 versus item 2) x 3 (interference condition: baseline, no-match, match) repeated measures ANOVA on the absolute error.

**Experiment 1:**

In Experiment 1, this analysis revealed three reliable main effects: Cue status showed a reliable main effect ($F$ (1, 47) = 58.84 $p < 0.001$; Figure S2A – blue versus orange bars). Serial position showed a significant main effect ($F$ (1,47) = 8.36, $p = 0.006$; Figure S2A – left versus right panel), with accuracy being higher slightly higher for the first encoded compared to the second encoded item. There was also a significant effect of interference ($F$ (1.48; 69.61) = 103.82, $p < 0.001$), showing that the interference manipulation worked across conditions. However, for interaction effects, only the presentation order x interference interaction reached significance ($F$ (1.76, 82.54) = 4.73; $p = 0.015$). All interactions involving the cue factor were non-significant (Cue × Position: $F < 1$, $p = .637$; Cue × Interference: $F < 1$, $p = .589$; Cue × Position × Interference: $F = 0.12$, $p = .868$) (For the full results table see Table 7). Thus, the cue benefit was strictly additive with both serial position and interference condition—the blue bars were consistently below the orange bars in every panel of Figure S2, but their relative spacing did not vary systematically across conditions.

To complement the frequentist analysis, we repeated the 2 (cue status) × 2 (presentation order) × 3 (interference condition) design with a Bayesian repeated-measures ANOVA in JASP. Model comparison converged on exactly the pattern described above. The model containing the three main effects and the presentation order × interference interaction received the highest posterior probability, $P$(M | data) = .52, and overwhelming evidence relative to the null model ($BF_{10} > 100$). Adding any interaction that involved the cue status reduced support: for example, including cue status × presentation order dropped the posterior probability to .046 ($BF_{10} > 100$) and including cue × interference to .017 ($BF_{10} > 100$). Across all models, inclusion Bayes factors indicated decisive evidence for retaining the cue main effect (BF_incl > 100) but evidence against retaining either cue status × presentation order or cue status × interference condition (both BF_incl < 0.5) (For the full table see Table 8)**.** This provided further evidence that cue manipulation yielded a robust, additive accuracy gain that did not interact with presentation order or with interference manipulation.

***Supplementary Table 7***

Full ANOVA table for the frequentist analysis of Experiment 1.

| **Effect** | **DF** | **MSE** | **F** | **p-value** |
|---|---|---|---|---|
| Cue status | 1,47 | 43.09 | 59.84 | <0.001 |
| Presentation order | 1,47 | 35.12 | 8.36 | 0.006 |
| Interference condition | 1.48,69.61 | 25.54 | 103.82 | <0.001 |
| Cue status * Presentation order | 1,47 | 14.38 | 0.23 | 0.637 |
| Cue status * Interference condition | 1.68,78.95 | 13.53 | 0.48 | 0.589 |
| Presentation order * Interference condition | 1.76,82.54 | 12.71 | 4.73 | 0.015 |
| Cue status * Presentation Order * Interference condition | 1.83,86.20 | 8.12 | 0.12 | 0.868 |

***Supplementary Table 8***

Full ANOVA table of Bayesian analysis of Experiment 1. Table ordered from best to worst model.

| **Model** | **P(M)** | **BF(M\|data)** | **$BF_M$** | **$BF_{10}$** | **Error (%)** |
|---|---|---|---|---|---|
| Null model (incl. subject + random slope | 0.053 | $5.732\times10^{-31}$ | $1.032\times10^{-29}$ | 1.000 | |
| Cue status + Presentation order + Interference condition + Presentation order * Interference condition | 0.053 | 0.517 | 19.249 | $9.015\times10^{+29}$ | 3.720 |
| Cue status + Presentation order + Interference condition | 0.053 | 0.195 | 4.361 | $3.403\times10^{+29}$ | 4.040 |
| Cue status + Presentation order + Interference condition + Cue status * Presentation order + Presentation order * Interference condition | 0.053 | 0.122 | 2.497 | $2.126\times10^{+29}$ | 7.451 |
| Cue status + Presentation order + Interference condition + Cue status * Interference condition + Presentation order * Interference condition | 0.053 | 0.058 | 1.100 | $1.005\times10^{+29}$ | 19.786 |

| | | | | | |
|---|---|---|---|---|---|
| Cue status + Presentation order + Interference condition + Cue status * Presentation Order | 0.053 | 0.046 | 0.864 | $7.988\times10^{+28}$ | 9.374 |
| Cue status + Interference condition | 0.053 | 0.029 | 0.533 | $5.017\times10^{+28}$ | 4.621 |
| Cue status + Presentation order + Interference condition + Cue status * Interference condition | 0.053 | 0.017 | 0.316 | $3.007\times10^{+28}$ | 4.927 |
| Cue status + Presentation order + Interference condition + Cue status * Presentation order + Cue status * Interference condition + Presentation order * Interference condition | 0.053 | 0.010 | 0.183 | $1.756\times10^{+28}$ | 4.580 |
| Cue status + Presentation order + Interference condition + Cue status * Presentation order + Cue status * Interference condition | 0.053 | 0.003 | 0.060 | $5.760\times10^{+27}$ | 3.834 |
| Cue status + Interference condition + Cue status * Interference condition | 0.053 | 0.003 | 0.049 | $4.706\times10^{+27}$ | 6.790 |
| Cue status + Presentation order + Interference condition + Cue status * Presentation order + Cue status * Interference condition + Presentation order * Interference condition + Cue status * Presentation order * Interference condition | 0.053 | $8.863\times10^{-4}$ | 0.016 | $1.546\times10^{+27}$ | 10.670 |
| Presentation order + Interference condition + Presentation order * Interference condition | 0.053 | $3.147\times10^{-8}$ | $5.664\times10^{-7}$ | $5.489\times10^{+22}$ | 4.031 |
| Presentation order + Interference condition | 0.053 | $1.122\times10^{-8}$ | $2.020\times10^{-7}$ | $1.958\times10^{+22}$ | 3.622 |
| Interference condition | 0.053 | $1.745\times10^{-9}$ | $3.141\times10^{-8}$ | $3.044\times10^{+21}$ | 4.654 |
| Cue status + Presentation order | 0.053 | $6.763\times10^{-23}$ | $1.217\times10^{-21}$ | $1.180\times10^{+8}$ | 4.619 |
| Cue status + Presentation order + Cue status * Presentation order | 0.053 | $1.302\times10^{-23}$ | $2.344\times10^{-22}$ | $2.272\times10^{+7}$ | 4.045 |
| Cue status | 0.053 | $9.493\times10^{-24}$ | $1.709\times10^{-22}$ | $1.656\times10^{+7}$ | 3.265 |
| Presentation order | 0.053 | $3.782\times10^{-30}$ | $6.808\times10^{-29}$ | 6.599 | 3.709 |

*Analysis of Effects (across all models)*

| Effects | P(incl) | P(excl) | P(incl\|data) | P(excl\|data) | $BF_{incl}$ |
|---|---|---|---|---|---|
| Cue status | 0.737 | 0.263 | 1.000 | $4.443\times10^{-8}$ | $8.038\times10^{+6}$ |
| Presentation order | 0.737 | 0.263 | 0.969 | 0.031 | 10.996 |
| Interference condition | 0.737 | 0.263 | 1.000 | $6.661\times10^{-16}$ | $5.361\times10^{+14}$ |
| Cue status * Presentation order | 0.316 | 0.684 | 0.182 | 0.818 | 0.482 |
| Cue status * Interference condition | 0.316 | 0.684 | 0.092 | 0.908 | 0.219 |
| Presentation order * Interference condition | 0.316 | 0.684 | 0.707 | 0.293 | 5.233 |
| Cue status * Presentation order * Interference condition | 0.053 | 0.947 | $8.863\text{x}10^{-4}$ | 0.999 | 0.016 |

**Experiment 2:**

For Experiment 2 we repeated the same analysis as above, and for this dataset we only found a main effect of Cue status ($F$ (1,40) = 31.22, $p$ < 0.001; Figure S2B – blue versus orange bars) and Interference condition ($F$ (1.60,67.70) = 32.87, $p$ < 0.001). All other main effects or interaction effects were not significant (see Supplementary table 9). No effect of presentation order is in line with the analyses above, and we found no effect of interference either, which is further in line with the analyses shown in the main text.

Nevertheless, to complement the frequentist analyses, we also repeated the Bayesian repeated-measures ANOVA. The model which contained the main effect of cue status and interference condition performed best (Posterior probability ($P$(M|data) = 0.517) and provided strong evidence relative to the null model ($BF_{10}$ > 100; for full table see Supplementary table 10). Across all models, inclusion Bayes factors for retaining both the cue status and interference condition main effects were the only factors with decisive evidence (BF_incl > 100).

***Supplementary Table 9***

Full ANOVA table for the frequentist analysis of Experiment 2.

| Effect | DF | MSE | F | p-value |
|---|---|---|---|---|
| Cue status | 1,40 | 15.14 | 31.22 | <0.001 |
| Presentation order | 1,40 | 26.01 | 2.62 | 0.113 |
| Interference condition | 1.69,67.70 | 10.55 | 32.87 | <0.001 |
| Cue status * Presentation order | 1,40 | 10.00 | 0.02 | 0.920 |
| Cue status * Interference condition | 1.92,76.67 | 6.48 | 0.20 | 0.812 |
| Presentation order * Interference condition | 1.97,78.83 | 7.42 | 0.70 | 0.496 |
| Cue status * Presentation Order * Interference condition | 1.98,79.38 | 10.22 | 0.35 | 0.708 |

***Supplementary Table 10***

Full ANOVA table of Bayesian analysis of Experiment 2. Table ordered from best to worst model.

| Model | P(M) | BF(M\|data) | $BF_M$ | $BF_{10}$ | Error (%) |
|---|---|---|---|---|---|
| Null model (incl. subject + random slope | 0.053 | $1.419\times10^{-12}$ | $2.554\times10^{-11}$ | 1.000 | |
| Cue status + Interference condition | 0.053 | 0.517 | 19.282 | $3.645\times10^{+11}$ | 2.159 |
| Cue status + Presentation order + Interference condition | 0.053 | 0.323 | 8.607 | $2.280\times10^{+11}$ | 2.489 |
| Cue status + Presentation order + Interference condition + Cue status * Presentation order | 0.053 | 0.060 | 1.140 | $4.197\times10^{+10}$ | 3.381 |
| Cue status + Interference condition + Cue status * Interference condition | 0.053 | 0.034 | 0.630 | $2.385\times10^{+10}$ | 4.121 |
| Cue status + Presentation order + Interference condition + Presentation order * Interference condition | 0.053 | 0.032 | 0.591 | $2.241\times10^{+10}$ | 2.613 |
| Cue status + Presentation order + Interference condition + Cue status * Interference condition | 0.053 | 0.022 | 0.397 | $1.519\times10^{+10}$ | 3.458 |
| Cue status + Presentation order + Interference condition + Cue status * Presentation order + Presentation order * Interference condition | 0.053 | 0.006 | 0.106 | $4.139\times10^{+9}$ | 4.071 |
| Cue status + Presentation order + Interference condition + Cue status * Presentation order + Cue status * Interference condition | 0.053 | 0.004 | 0.070 | $2.734\times10^{+9}$ | 3.635 |
| Cue status + Presentation order + Interference condition + Cue status * Interference condition + Presentation order * Interference condition | 0.053 | 0.002 | 0.035 | $1.366\times10^{+9}$ | 2.474 |
| Cue status + Presentation order + Interference condition + Cue status * Presentation order + Cue status * Interference condition + Presentation order * Interference condition | 0.053 | $7.196\times10^{-4}$ | 0.013 | $5.072\times10^{+8}$ | 49.777 |
| Interference condition | 0.053 | $8.272\times10^{-5}$ | 0.001 | $5.830\times10^{+7}$ | 1.193 |

| | | | | | |
|---|---|---|---|---|---|
| Presentation order + Interference condition | 0.053 | $5.679\times10^{-5}$ | 0.001 | $4.003\times10^{+7}$ | 6.053 |
| Cue status + Presentation order + Interference condition + Cue status * Presentation order + Cue status * Interference condition + Presentation order * Interference condition + Cue status * Presentation order * Interference condition | 0.053 | $4.441\times10^{-5}$ | $7.995\times10^{-4}$ | $3.130\times10^{+7}$ | 2.998 |
| Presentation order + Interference condition + Presentation order * Interference condition | 0.053 | $5.153\times10^{-6}$ | $9.275\times10^{-5}$ | $3.632\times10^{+6}$ | 2.696 |
| Cue status | 0.053 | $8.392\times10^{-9}$ | $1.511\times10^{-7}$ | 5915.018 | 1.358 |
| Cue status + Presentation Order | 0.053 | $5.137\times10^{-9}$ | $9.246\times10^{-8}$ | 3620.337 | 2.509 |
| Cue status + Presentation order + Cue status * Presentation order | 0.053 | $1.041\times10^{-9}$ | $1.873\times10^{-8}$ | 733.558 | 6.822 |
| Presentation order | 0.053 | $8.602\times10^{-13}$ | $1.548\times10^{-11}$ | 0.606 | 1.310 |

*Analysis of Effects (across all models)*

| **Effects** | **P(incl)** | **P(excl)** | **P(incl\|data)** | **P(excl\|data)** | **$BF_{incl}$** |
|---|---|---|---|---|---|
| Cue status | 0.737 | 0.263 | 1.000 | $1.447\times10^{-4}$ | 2468.398 |
| Presentation order | 0.737 | 0.263 | 0.449 | 0.551 | 0.291 |
| Interference condition | 0.737 | 0.263 | 1.00 | $1.457\times10^{-8}$ | $2.451\times10^{+7}$ |
| Cue status * Presentation order | 0.316 | 0.684 | 0.060 | 0.930 | 0.163 |
| Cue status * Interference condition | 0.316 | 0.684 | 0.062 | 0.938 | 0.143 |
| Presentation order * Interference condition | 0.316 | 0.684 | 0.040 | 0.960 | 0.091 |
| Cue status * Presentation order * Interference condition | 0.053 | 0.947 | $4.441\times10^{-5}$ | 1.000 | $7.995\times10^{-4}$ |

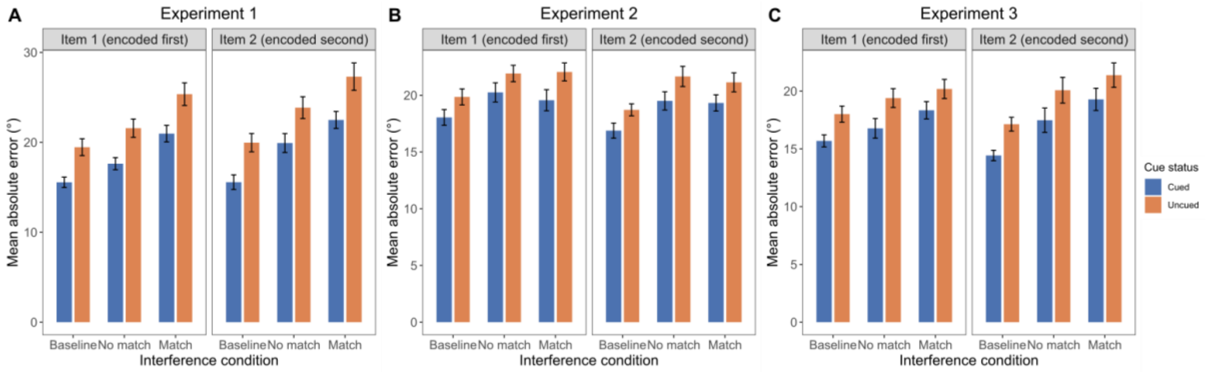


*Figure 2:* Absolute error across the three interference conditions in Experiment 1 (**A**), Experiment 2 (**B**) and Experiment 3 (**C**) (baseline, no match and match) split for item presentation (item 1 – left panel; item 2 – right panel), depending on if the item was cued (prioritised; in blue) or uncued (deprioritised; in orange). Boxplot shows the mean, and error bars indicate the SEM.

**Experiment 3:**

In Experiment 3, we found a main effect of cue status ($F$ (1, 43) = 41.99 $p$ < 0.001; Figure S2C – blue versus orange bars) and interference condition ($F$ (1.65,70.93) = 12.05 $p$ < 0.001) but not for presentation order ($F$ (1,43) = 0.28; see Table 11 for all values). As for Experiment 1, we found an interaction effect of presentation order and interference condition ($F$ (1.99,85.61) = 4.83, $p$ = 0.01) but no other interaction effects were significant (Table 11 for full ANOVA values). Thus, we confirmed our conclusion from Experiment 1 that any cue benefit was purely additive; and their spacing did not vary systematically across conditions.

We again repeated the analyses as a Bayesian repeated-measures ANOVA. Model comparisons revealed a model with both cue status and interference condition as main effects as the best model ($P$(M | data) = .4; $BF_{10}$ > 100). Looking at the effects, we found decisive evidence for an effect of cue status and interference condition ($BF_{incl}$ > 100 for both); while we only found anecdotal evidence for an effect of the interaction term which we found in our frequentist analyses (Presentation order * Interference condition; $BF_{incl}$ = 1.663).

***Supplementary Table 11***

Full ANOVA table for the frequentist analysis of Experiment 3.

| **Effect** | **DF** | **MSE** | **F** | **p-value** |
|---|---|---|---|---|
| Cue status | 1,43 | 17.47 | 41.99 | <0.001 |
| Presentation order | 1,43 | 24.80 | 0.28 | 0.597 |
| Interference condition | 1.65,70.93 | 54.51 | 12.05 | <0.001 |
| Cue status * Presentation order | 1,43 | 7.97 | 0.17 | 0.681 |
| Cue status * Interference condition | 1.97,84.52 | 10.60 | 0.50 | 0.605 |
| Presentation order * Interference condition | 1.99,85.61 | 11.91 | 4.82 | 0.010 |
| Cue status * Presentation Order * Interference condition | 1.80,77.50 | 8.12 | 0.07 | 0.92 |

***Supplementary Table 12***

Full ANOVA table of Bayesian analysis of Experiment 2. Table ordered from best to worst model.

| Model | P(M) | BF(M\|data) | $BF_M$ | $BF_{10}$ | Error (%) |
|---|---|---|---|---|---|
| Null model (incl. subject + random slope | 0.053 | $3.357\times10^{-9}$ | $6.042\times10^{-8}$ | 1.000 | |
| Cue status + Interference condition | 0.053 | 0.404 | 12.183 | $1.202\times10^{+8}$ | 4.304 |
| Cue status + Presentation order + Interference condition + Presentation order * Interference condition | 0.053 | 0.335 | 9.069 | $9.981\times10^{+7}$ | 6.196 |
| Cue status + Presentation order + Interference condition | 0.053 | 0.092 | 1.823 | $2.740\times10^{+7}$ | 4.789 |
| Cue status + Presentation order + Interference condition + Cue status * Presentation order + Presentation order * Interference condition | 0.053 | 0.058 | 1.100 | $1.716\times10^{+7}$ | 6.139 |
| Cue status + Interference condition + Cue status * Interference condition | 0.053 | 0.044 | 0.830 | $1.313\times10^{+7}$ | 6.510 |
| Cue status + Presentation order + Interference condition + Cue status * Interference condition + Presentation order * Interference condition | 0.053 | 0.035 | 0.652 | $1.041\times10^{+7}$ | 8.161 |
| Cue status + Presentation order + Interference condition + Cue status * Presentation order | 0.053 | 0.016 | 0.285 | $4.638\times10^{+6}$ | 4.486 |
| Cue status + Presentation order + Interference condition + Cue status * Interference condition | 0.053 | 0.008 | 0.149 | $2.450\times10^{+6}$ | 4.153 |
| Cue status + Presentation order + Interference condition + Cue status * Presentation order + Cue status * Interference condition + Presentation order * Interference condition | 0.053 | 0.006 | 0.112 | $1.838\times10^{+6}$ | 6.471 |
| Cue status + Presentation order + Interference condition + Cue status * Presentation order + Cue status * Interference condition | 0.053 | 0.002 | 0.030 | 489890.659 | 5.845 |
| Cue status | 0.053 | $5.200\times10^{-4}$ | 0.009 | 154902.197 | 8.503 |
| Cue status + Presentation order + Interference condition + Cue status * Presentation order + Cue | 0.053 | $4.708\times10^{-4}$ | 0.008 | 140250.310 | 8.842 |

| | | | | | |
|---|---|---|---|---|---|
| status * Interference condition + Presentation order * Interference condition + Cue status * Presentation order * Interference condition | | | | | |
| Cue status + Presentation order | 0.053 | $9.531 \times 10^{-5}$ | 0.002 | 28392.139 | 4.501 |
| Cue status + Presentation order + Cue status * Presentation order | 0.053 | $1.662 \times 10^{-5}$ | $2.992 \times 10^{-4}$ | 4950.981 | 4.797 |
| Interference condition | 0.053 | $2.991 \times 10^{-6}$ | $5.384 \times 10^{-5}$ | 890.981 | 3.975 |
| Presentation order + Interference condition + Presentation order * Interference condition | 0.053 | $2.341 \times 10^{-6}$ | $4.214 \times 10^{-5}$ | 697.351 | 6.014 |
| Presentation order + Interference condition | 0.053 | $6.301 \times 10^{-7}$ | $1.134 \times 10^{-5}$ | 187.722 | 5.001 |
| Presentation order | 0.053 | $7.247 \times 10^{-10}$ | $1.305 \times 10^{-8}$ | 0.216 | 5.892 |

*Analysis of Effects (across all models)*

| **Effects** | **P(incl)** | **P(excl)** | **P(incl\|data)** | **P(excl\|data)** | $\mathbf{BF_{incl}}$ |
|---|---|---|---|---|---|
| Cue status | 0.737 | 0.263 | 1.000 | $5.966 \times 10^{-6}$ | 59862.946 |
| Presentation order | 0.737 | 0.263 | 0.552 | 0.448 | 0.440 |
| Interference condition | 0.737 | 0.263 | 0.999 | $6.319 \times 10^{-4}$ | 564.823 |
| Cue status * Presentation order | 0.316 | 0.684 | 0.081 | 0.919 | 0.192 |
| Cue status * Interference condition | 0.316 | 0.684 | 0.096 | 0.904 | 0.229 |
| Presentation order * Interference condition | 0.316 | 0.684 | 0.434 | 0.566 | 1.663 |
| Cue status * Presentation order * Interference condition | 0.053 | 0.947 | $4.708 \times 10^{-4}$ | 1.000 | 0.008 |

Taken together, these analyses showed that the block-wise cue reliably improve memory performance in Experiment 1-Experiment 3, over and above any effects of presentation order or interference condition. The cue benefit was additive rather than interactive with these factors, indicating that performance is systematically better for the cued item. In combination with the results of Experiment 4 in the main text, where cue status and response order were explicitly decoupled, this supports the interpretation that the block cue effectively prioritised one item in working memory, rather than merely reflecting output order or recency effects.

**3. Additional analyses for Experiment 4**

***Supplementary Table 13***

Full ANOVA table of 2 (Cue order) x 2 (Probe order) analysis of Experiment 4. Table ordered from best to worst model.

| **Model** | **P(M)** | **BF(M\|data)** | **$BF_M$** | **$BF_{10}$** | **Error (%)** |
|---|---|---|---|---|---|
| Null model (incl. subject + random slope | 0.2 | $2.247\times10^{-6}$ | $8.986\times10^{-6}$ | 1.000 | |
| Cue condition + Probe order | 0.2 | 0.746 | 11.751 | 332087.726 | 4.595 |
| Cue condition + Probe order + Cue condition * Probe order | 0.2 | 0.140 | 0.649 | 62138.073 | 2.652 |
| Probe order | 0.2 | 0.114 | 0.516 | 50894.860 | 1.595 |
| Cue condition | 0.2 | $1.232\times10^{-5}$ | $4.928\times10^{-5}$ | 5.484 | 2.149 |

*Analysis of Effects (across all models)*

| **Effects** | **P(incl)** | **P(excl)** | **P(incl\|data)** | **P(excl\|data)** | **$BF_{incl}$** |
|---|---|---|---|---|---|
| Cue status | 0.6 | 0.4 | 0.886 | 0.114 | 5.164 |
| Probe order | 0.6 | 0.4 | 1.000 | $1.457\times10^{-5}$ | 45768.115 |
| Cue status * Probe order | 0.2 | 0.8 | 0.140 | 0.860 | 0.649 |
